# Calculation of ATP production rates using the Seahorse XF Analyzer

**DOI:** 10.1101/2022.04.16.488523

**Authors:** Brandon R. Desousa, Kristen K.O. Kim, Anthony E. Jones, Andréa B. Ball, Wei Y. Hsieh, Pamela Swain, Danielle H. Morrow, Alexandra J. Brownstein, David A. Ferrick, Orian S. Shirihai, Andrew Neilson, David A. Nathanson, George W. Rogers, Brian P. Dranka, Anne N. Murphy, Charles Affourtit, Steven J. Bensinger, Linsey Stiles, Natalia Romero, Ajit S. Divakaruni

## Abstract

Oxidative phosphorylation and glycolysis are the dominant ATP-generating pathways in mammalian metabolism. The balance between these two pathways is often shifted to execute cell-specific functions in response to stimuli that promote activation, proliferation, or differentiation. However, measurement of these metabolic switches has remained mostly qualitative, making it difficult to discriminate between healthy, physiological changes in energy transduction or compensatory responses due to metabolic dysfunction. We therefore developed a broadly applicable method to calculate ATP production rates from oxidative phosphorylation and glycolysis using Seahorse XF Analyzer data. We quantified the bioenergetic changes observed during macrophage polarization as well as cancer cell adaptation to *in vitro* culture conditions. Additionally, we detected substantive changes in ATP utilization upon neuronal depolarization and T cell receptor activation that are not evident from steady-state ATP measurements. This method generates a single readout that allows the direct comparison of ATP produced from oxidative phosphorylation and glycolysis in live cells. Additionally, the manuscript provides a framework for tailoring the calculations to specific cell systems or experimental conditions.

## INTRODUCTION

ATP is critical for cellular energy transduction. Essential cellular functions such as maintenance of ion homeostasis, macromolecule biosynthesis, motility, and autophagy involve reactions coupled to ATP hydrolysis to support these otherwise energetically unfavorable processes. The two dominant ATP-producing pathways in mammalian cells are glycolysis and oxidative phosphorylation (Chandel, 2015). Glycolysis involves the cytoplasmic breakdown of one molecule glucose into two molecules of pyruvate, generating two molecules of ATP in the process. The fermentation of pyruvate to lactate and subsequent lactate efflux enables continuous glycolytic turnover. Pyruvate and other carbon-rich energy substrates such as fatty acids and amino acids can also be taken up by mitochondria and completely oxidized during oxidative phosphorylation. The chemical energy in these substrates is converted to potential energy by the mitochondrial respiratory chain, and this reservoir of potential energy is used to drive ATP synthesis (Rich & Maréchal, 2010; Nicholls & Ferguson, 2013).

It is now clear that redistributing the balance between oxidative phosphorylation and glycolysis can characterize – and in some cases determine – cell function and fate (Lunt & Vander Heiden, 2011; DeBerardinis & Chandel, 2020; Sullivan *et al*, 2018; Shin *et al*, 2020). Glycolysis and offshoot pathways play essential roles in generating precursors for macromolecule biosynthesis(Lunt & Vander Heiden, 2011), supporting one-carbon metabolism (Ducker & Rabinowitz, 2017), and establishing cytoplasmic reducing power (Lewis *et al*, 2014). Biological processes such as stem cell pluripotency(Gu *et al*, 2016), oncogenesis (DeBerardinis & Chandel, 2020), growth factor signaling (Pavlova & Thompson, 2016), and immune cell activation (Van den Bossche *et al*, 2017) are therefore frequently associated with a preferential shift towards glycolysis. Conversely, terminal differentiation is often accompanied by an increased capacity for oxidative phosphorylation and the complete oxidation of glucose and fatty acids (Chen *et al*, 2008; Wilson-Fritch *et al*, 2003; Divakaruni *et al*, 2014; Moyes *et al*, 1997; Hazim *et al*, 2022).

Despite the importance of cellular ATP production rates and the distribution between glycolysis and oxidative phosphorylation, common measurement approaches are often limiting. Snapshot measurements of cellular ATP content are routinely used to characterize changes in energy metabolism, but these are generally informative only in extreme cases of energetic dysfunction such as ischemia (Brand & Nicholls, 2011). During physiological cell activation, cells can readily increase the rate of ATP production to match the increased rate of consumption without appreciably changing steady-state ATP levels. A far more useful metric is the ATP:ADP ratio, reflecting the free energy change associated with ATP hydrolysis into ADP and inorganic phosphate. However, biochemical or mass spectrometry-based approaches to measure ATP:ADP are limited given that they are destructive measurements conducted on cell lysates and unable to measure real-time changes and are prone to experimental noise that masks subtle but meaningful differences. Kinetic measurements of energy status are therefore extraordinarily specialized, requiring transfection of a genetically engineered ATP:ADP sensor and an expert understanding of the caveats when using molecular probes (Tantama & Yellen, 2014; Koveal *et al*, 2020). Importantly, all these approaches lack direct information about whether the measured ATP is derived from oxidative phosphorylation or glycolysis.

Fluorescence-based, multi-well technologies such as Seahorse XF Analysis can characterize these bioenergetic shifts, as simultaneous measurements of oxygen consumption rates (OCR) and extracellular acidification rates (ECAR) can gauge the relative balance between oxidative phosphorylation and glycolysis (Pelletier *et al*, 2014; Divakaruni & Jastroch, 2022) (i.e. OCR:ECAR ratio). However, precise quantification of the total rate of ATP production and the balance between these two pathways is not straightforward. Direct comparison is precluded by differences in unit scale between OCR (pmol O_2_/min) and ECAR (mpH/min). Direct, quantitative comparison between the two pathways is further confounded by the differential ATP yields from each pathway: for oxidation of one molecule of glucose, the maximum glycolytic yield is 2 ATP as opposed to 33.45 ATP for oxidative phosphorylation (Mookerjee *et al*, 2017). As such, a large increase in acidification of the extracellular medium may not necessarily indicate a rebalancing of ATP production if it is matched with even a marginal increase in oxygen consumption.

Additionally, ECAR measurements can be influenced by cellular processes other than lactate efflux that also result in a net acidification of the extracellular medium. Indeed, the main component of ECAR in two-dimensional cell culture systems is often glycolysis and fermentation (McConnell *et al*, 1992), reflecting the uptake of uncharged glucose and the release of anionic lactate. However, other acidifying reactions can contribute to the measured ECAR. For example, CO_2_ evolution during oxidative metabolism (e.g. dehydrogenases of the TCA cycle or the pentose phosphate pathway) acidifies the medium after hydration and generation of bicarbonate (CO_2_ + H_2_O → H_2_CO_3_ ® H^+^ + HCO^3-^). Additionally, specific cell types can release appreciable amounts of organic acids such as pyruvate (Hong *et al*, 2016), glutamate (Divakaruni *et al*, 2017), fatty acids (Thompson *et al*, 2010)that may be reflected in ECAR measurements. As such, the extracellular acidification rate cannot itself quantify lactate efflux (Newell *et al*, 1993; Gillies *et al*, 2008; Divakaruni *et al*, 2014).

We therefore developed a method to transform OCR and ECAR into rates of ATP production from oxidative phosphorylation and glycolysis. Although measurements of OCR and ECAR have become a central cell biology technique, the qualitative nature of the analysis makes it difficult to discriminate between healthy, physiological shifts in bioenergetic pathways or compensatory responses due to mitochondrial dysfunction. This method detailed here provides a solution to this challenge in a single, live cell readout estimating the total rate of cellular ATP production as well as its distribution between oxidative phosphorylation and glycolysis. Model parameters are formed with data from over 20 primary and immortalized cell lines, and the analysis detects substantive changes not evident from steady-state ATP measurements. Although a broadly similar approach heavily reliant on reductionist systems has been previous used, the values obtained from this largely empirical strategy sharply contrast previous work (Mookerjee *et al*, 2015).

## MATERIALS AND METHODS

### Chemicals

All chemicals were purchased from Sigma-Aldrich unless otherwise stated.

### Animals and primary cells

All animal procedures were performed in accordance with the NIH Guide for the Care and Use of Laboratory Animals, and approved by the UCLA Animal Research Committee (ARC-2020-027).

#### Murine bone marrow-derived macrophages (BMDMs)

BMDMs were cultured as previously described (Hsieh *et al*, 2020). Briefly, macrophages were isolated from femurs of male C57BL/6J mice (Jackson 000664) and maintained at 37°C in a humidified 5% CO_2_ incubator. Cells were treated with 3 mLs RBC lysis buffer (Sigma-Aldrich) to remove red blood cells for 5 minutes, centrifuged at 386*g* for 5 minutes, and resuspended in BMDM culture medium. Culture medium consisted of high-glucose DMEM (Gibco 11965) supplemented with 10% heat-inactivated fetal bovine serum (FBS), 2 mM L-glutamine, 100 U/ml penicillin, 100 μg/mL streptomycin, 500 µM sodium pyruvate, and 5% (v/v) conditioned medium containing macrophage colony stimulating factor (M-CSF) produced by CMG cells to induce differentiation to BMDMs. BMDMs were allowed to differentiate for 6 days, changing medium after four days, and then plated for experiments.

#### Rat cortical neurons

Cortical neurons were generated from embryonic day 18 Sprague-Dawley rats as described previously (Kushnareva *et al*, 2005) and maintained at 37°C in a humidified 5% CO_2_ incubator. Cells were plated onto poly-d-lysine-coated wells of black-walled 96-well plates for enzymatic assays or Seahorse XF96 plates. Cells for 96-well plate-based assays were seeded at 2.5×10^4^ cells/well and maintained in maintenance medium composed of 200 μL Neurobasal (Thermo Fisher Scientific) medium supplemented with 1X B27 serum-free supplement (Thermo Fisher Scientific), 2 mM GlutaMAX (Thermo Fisher Scientific), 100 U/ml penicillin, and 100 μg/mL streptomycin. Half the medium was replaced every 3–4 d. Experiments in 96-well plates were conducted at day in vitro (DIV) 13–17.

#### Human T cells

Frozen stocks of human naïve CD4^+^ T cells were purchased from Lonza. Cells were thawed and permitted to recover in RPMI 1640 (Gibco, #11875-085) supplemented with 10%(v/v) FBS at 37°C, 5% CO_2_ for 18-24 hr. After recovery, cells were spun at 250*g* for 10 min and resuspended to a final density of 4 x 10^6^ cells/mL in Agilent Seahorse XF RPMI Medium pH 7.4 (# 103576-100) supplemented with 10 mM glucose, 2 mM glutamine and 1 mM pyruvate (XF Assay Media). 50 µL of the naïve T-cell suspension (2 x 10^5^ cells) were plated in each well of a poly-D-lysine-coated Seahorse XF96 microplate. Plates were spun at 200*g* for 1 min to adhere cells to the plate. 130 µL of warmed XF Assay Media was then added to each well, and plates were either run in the XF Analyzer or used to measure intracellular ATP concentration with the Cell-Titer Glo Assay (Promega). ATP measurements were taken before and after activation with magnetic Dynabeads coated with anti-CD3/CD28 antibodies (Thermofisher, 4:1 beads/cell).

#### Additional primary cells preparations

Neonatal rat ventricular myocytes (Rubio *et al*, 2009) and rat cortical astrocytes (Kim & Magrané, 2011) were isolated as previously described. Murine splenocytes were isolated from dissected spleens from C57BL/6 mice. Tissue was mechanically dissociated in HBSS and passed through a 40 µm strainer. Cells were washed and resuspended in red blood cells lysis buffer (Stemcell # 07850) and incubated on ice for 10 min. Cell were washed again in HBSS, passed through a 70 µm filter, counted, and used immediately. For Seahorse XF experiments cells were resuspended in XF Assay Media and seeded at 1.5 x 10^5^ cells/well.

#### Mitochondrial Isolation

Hearts from C57BL/6 mice were drained of blood and homogenized with a hand-held Polytron followed by a Potter-Elvehjem Teflon-on-glass tissue grinder in ice-cold MSHE buffer [70 mM sucrose, 210 mM mannitol, 5 mM HEPES, 1 mM EGTA, and 0.5% (w/v) fatty acid-free BSA](Kubli *et al*, 2013). The homogenate was centrifuged at 500*g* for 10 minutes at 4°C. The supernatant was then collected and centrifuged at 12,000 *g* for 10 minutes at 4°C to obtain a mitochondrial pellet. The pellet was washed in MSHE and centrifuged at 12,000 *g* for 10 minutes at 4°C, and this wash step was repeated. Finally, the pellet was resuspended in a small volume (∼20 μL) of MSHE, and the protein concentration was determined by Bradford assay.

### Cultured cells

All cell lines were obtained from American Type Culture Collection (ATCC) and cultured as suggested by the supplier except when otherwise notified. Each formulation of medium was supplemented with 10% (v/v) fetal bovine serum (FBS), 2 mM GlutaMAX, 100 U/mL penicillin, and 100 μg/mL streptomycin unless otherwise specified. All cells were maintained at 37°C in a humidified 5% CO_2_ incubator. Cell lines were maintained in the medium listed parenthetically: A549 [DMEM/F12 (Gibco 11330); HepG2 [MEM (Gibco 11095)]; HCT116 [RPMI (Gibco 11875), C2C12 [DMEM (Gibco 11960 + 2 mM glutamine + 1 mM pyruvate)], A431 [DMEM (Gibco 11960) + 2 mM glutamine + 1 mM pyruvate), BAEC [DMEM (Gibco 11054)], BT474 [ATCC Hybri-Care Cat#46-x], H460 [RPMI (Gibco 11870), 1 mM HEPES, 4 mM glutamine, 5% FBS), HUVEC [EGM Endothelial Cell Growth Medium (Lonza CC-3124)], Jurkat,[RPMI (Gibco A10491-01)], MCF7 [RPMI (Gibco 21870 + 2 mM glutamine + 1 mM pyruvate)], MCF-10A [MEBM supplemented with Kit CC-3150 (Lonza) and 100 ng/mL cholera toxin (Sigma)], MDA-MB231 [RPMI (Gibco 21870 + 2 mM glutamine + 1 mM pyruvate)], PC12 [RPMI (ATCC 30-2001 + 2mM glutamine + 10% Horse Serum + 5% FBS], Raw 264.7 [DMEM (Gibco 11960) + 2 mM glutamine + 1 mM pyruvate].

#### 3T3-L1 pre-adipocytes and differentiation

3T3-L1 pre-adipocytes were maintained below ∼70% confluency in DMEM (Gibco 11965) supplemented with 10% (v/v) bovine calf serum (BCS), 100 U/mL penicillin, and 100 μg/mL streptomycin. For differentiation, cells were plated on Day -2 and were allowed to reach confluency over 48 hr. Differentiation was induced on Day 0 by addition of fresh medium with FBS replacing BCS and supplemented with the following: 0.5 mM IBMX, 0.25 μM dexamethasone, 1 μg/ml insulin, and 100 nM rosiglitazone. On Day 2, medium was replaced to DMEM + FBS supplemented with only 1 μg/ml insulin, and thereafter insulin was removed when replacing medium every two days.

#### Immortalized brown adipocytes

Immortalized human brown adipocytes were maintained as described previously (Panic *et al*, 2020). Briefly, cells were cultured in DMEM supplemented with 25 mM glucose, 10% (v/v) fetal bovine serum, 4 mM glutamine, 1 mM sodium pyruvate, 10 mM HEPES, and 100 U/ml penicillin with 100 µg/ml streptomycin. Cells were plated at 1×10^4^ cells/well and reached total confluency after approximately 24 hr. Medium was then changed to induction medium consisting of the growth medium as before supplemented with 1 µM rosiglitazone, 0.5 mM IBMX, 0.5 μM dexamethasone, 125µM Indomethacin, 1 nM T3, and 4 nM human insulin. After two days, medium was changed to growth medium supplemented with only rosiglitazone and insulin (concentrations as before), and this medium composition was used to change the fluid every other day for an additional 5-6 days.

### Seahorse XF Analysis

All oxygen consumption and extracellular acidification measurements were conducted using an Agilent Seahorse XF96 or XF^e^96 Analyzer. Experiments were conducted at 37°C and at pH 7.4 (intact cells) or 7.2 (isolated mitochondria). For experiments with cultured cells, only the inner 60 wells were used, and the outer rim was filled with 200 μL of PBS throughout the incubation to minimize variance in temperature and evaporative effects across the plate (Lundholt *et al*, 2003). All respiratory parameters were corrected for non-mitochondrial respiration and background signal from the instrument with addition of 200 nM rotenone and 1 μM antimycin A. Where appropriate, oligomycin was used at 2 μM unless otherwise specified, and FCCP concentrations were titrated to determine an optimal concentration for a given experiment. Unbuffered DMEM assay medium is composed of DMEM (Sigma #5030; pH 7.4) supplemented with 31.6 mM NaCl, 3 mg/L phenol red, and 5 mM HEPES unless otherwise indicated.

#### XF analysis primary cortical neurons

All experiments with primary cortical neurons were conducted in a custom, modified Neurobasal-type medium (Divakaruni *et al*, 2017) rather than DMEM to maintain appropriate osmolarity. Glutamine was omitted from the assay medium to minimize excitotoxic injury.

#### Acid injections and calculation of buffering power

Sequential injections of 62.5 nmol H^+^ (25 μL of 1.25 mM H_2_SO_4_) were made using the Seahorse XF injector ports into unbuffered DMEM assay medium (see above) supplemented with 8 mM glucose, 2 mM glutamine, 2 mM pyruvate and HEPES as indicated in the text. Values were corroborated with matched experiments using 2.5 mM HCl made from freshly purchased stocks no more than 2 months old. Calculations were made using a measurement microchamber volume of 2.28 μL. Step-by-step sample calculations for using the buffering power of the medium to convert ECAR (mpH/min) to pmol H^+^/min are provided in the Supplemental Worksheet.

#### Calculation of lactate:H^+^ and H^+^:O_2_

Rates of medium acidification without contribution from respiration were calculated as follows: DMEM assay medium was supplemented with 5 mM glucose and varying concentrations of 2-deoxyglucose (2-DG) and 5 mM iodoacetate (IA) where indicated in the figure. Medium was also supplemented with 200 nM rotenone (to block complex I), 1 μM antimycin A (to block complex III), and 2 μM oligomycin (to block ATP hydrolysis and prevent an energy crisis). Rates were calculated for 75 min, and the spent medium was immediately harvested from each plate and frozen at -80°C for later calculation of lactate efflux by enzymatic assay (Mookerjee *et al*, 2015).

Rates of medium acidification in the absence of glycolysis were calculated as follows: DMEM assay medium was supplemented with 5 mM pyruvate and 5 mM glutamine (no glucose was provided) along with 2 mM 2-DG and 50 μM IA (to block residual glycolysis and glycogenolysis). 500 U/mL carbonic anhydrase was added to ensure rapid CO_2_ hydration, though it was later determined to have a negligible effect on rates of acidification. Respiratory rates were titrated by addition of oligomycin, FCCP (500 nM or 1 μM), and rotenone with antimycin A. The medium was also supplemented with varying concentrations of both UK5099 and BPTES (3 nM - 3 μM for both) to further titrate the respiratory rate. Experiments with primary cortical neurons were conducted in custom medium lacking glutamine as described earlier.

#### Patient-derived glioblastoma cultures

Patient-derived high-grade gliomas were isolated, purified, and maintained as cell lines or implanted as orthotopic xenografts according to previously published protocols (Gosa *et al*, 2019). Cultured cells (*in vitro*) and tumor cells acutely purified from xenografts (*ex vivo*) were seeded onto Cell-Tak-coated XF96 plates (25 μg/well; Corning). Cells were plated at either 4.0 x 10^4^ or 1.0 x 10^5^ based on cell size and the initial oxygen consumption rate, and spun at 600*g* for 5 min to adhere cells to the well.

#### Cell activation assays

Cortical neurons were plated at 2.5×10^4^ cells/well and acutely activated with 1.5 μM veratridine added via the injector port of the instrument (Choi *et al*, 2009). Where indicated, cells were pretreated with 2 nM ouabain immediately prior to the assay. When activating the NMDA receptor, 100 μM NMDA was acutely given via the injector port to neurons with or without 15 min. pre-treatment of 10 μM MK-801. For these experiments, the custom assay medium was supplemented with 2 mM β-hydroxybutyrate as substrate, as pyruvate could not be used due to complications with measuring glutamate in the medium by a linked enzymatic assay to confirm depolarization and glutamate efflux (data not shown). For acute, T cell activation, murine anti-CD3/CD28 Dynabeads (Gibco) were added via the injector port as previously described (Gubser *et al*, 2013).

#### LPS-mediated activation of BMDMs

BMDMs were plated at 5.0×10^4^ cells/well for 48 hr. prior to cytokine treatment, which lasted for 24 hr. unless otherwise specified. Lipopolysaccharide (LPS) was purchased from PeproTech, Inc. (New Jersey, USA) and offered to cells at 50 ng/mL unless otherwise specified in the manuscript.

#### H^+^:O_2_ in isolated heart mitochondria

Isolated heart mitochondria were plated in XF96 plates (1.5 μg/well) and spun 2000*g* for 20 min to adhere mitochondria to the well. Assays were conducted in MAS medium ([220 mM mannitol, 70 mM sucrose, 10 mM KH_2_PO_4_, 5 mM MgCl_2_, 1 mM EGTA, and 2 mM HEPES (pH 7.2)] supplemented with 0.2% (w/v) fatty acid-free BSA. For all conditions, uncoupler-stimulated respiration was measured in the presence of 2 µM oligomycin and 4 µM FCCP to eliminate effects of ATP synthesis or hydrolysis on the acidification rate. Rates of respiration and acidification were measured under three conditions: (i) pyruvate/malate (P/M): 10 mM pyruvate with 1 mM malate and 2 mM dichloroacetate; (ii) pyruvate/malate with an inhibitor mix to block various enzymes to run a truncated TCA cycle (P/M/truncated): 10 mM pyruvate with 1 mM malate, 2 mM dichloroacetate, 60 μM fluorocitrate (to block isocitrate dehydrogenase), 2 mM malonate (to block succinate dehydrogenase), and 1 mM aminooxyacetate (to block transaminase activity), and (iii) succinate/rotenone (S/R): 10 mM succinate with 2 μM rotenone to consume oxygen without release of CO_2_ to acidify the experimental medium(Mookerjee *et al*, 2015).

### Normalization of Seahorse XF assays to total protein or cell number

Where indicated on the axis, Seahorse XF assays were normalized either to total protein or cell number determined by *post hoc* high-content imaging. When measuring total protein, cells were lysed in RIPA buffer [50 mM Tris, pH 7.4, 150 mM NaCl,1% (w/v) NP-40, 0.5% (w/v) sodium deoxycholate, and 0.1% (w/v) sodium dodecyl sulfate], boiled, and assayed using the bicinchoninic assay method (ThermoFisher).

When conducting *post hoc* cell counts, Seahorse XF plates were collected after assays, washed with PBS, and fixed overnight with 2% (w/v) paraformaldehyde in PBS. Cells were left fixed up to 2 weeks. On the day prior to counting, 10 ng/mL Hoescht 33342 (ThermoFisher) was added and incubated overnight at 4°C. Cells were counted using a PerkinElmer Operetta high-content, wide-field, fluorescence imaging system (ex./em. 361/497 nm) coupled to Columbus software (Operetta; PerkinElmer, Waltham, MA, USA).

### Enzymatic lactate measurements

Enzymatic lactate measurements were determined in the spent medium recovered after an Agilent Seahorse XF assay. Medium was brought to room temperature if previously frozen, and mixed 1:1 with a 2X assay solution of 40 U/mL lactate dehydrogenase (Sigma-Aldrich L3916), 1 M Tris (pH 9.8), 20 mM EDTA, 40 mM hydrazine (Sigma-Aldrich 309400), and 4 mM NAD^+^. The reaction velocity was given by NADH fluorescence (ex./em. 340/460) and measured using a Tecan Spark multimode plate reader (Mookerjee *et al*, 2015). Measurements were taken after 2-4 min (readings were taken every 30 sec. to determine when stable values were achieved), and calibrated against known lactate concentrations (Sigma-Aldrich L7022). Importantly, lactate standards were made in assay medium matching the experimental medium to account for any effects from millimolar concentrations of pyruvate in some experimental medium. Standard curves were calculated between 0.2 and 100.0 nmol lactate and modeled with a Michaelis-Menten curve using GraphPad Prism 5.0.

### ATP Content

Cellular ATP content was measured in lysed cells with the Cell Titer Glo assay (Promega) in 96-well plates according to the manufacturer’s specifications.

### Quantification and statistical analysis

All statistical parameters, including the number of replicates (n), can be found in the figure legends. Statistical analyses were performed using Graph Pad Prism 5 software for Mac OS X. Data are presented as the mean ± SEM unless otherwise specified. Individual pairwise comparisons were performed using two-tailed Student’s t-test. For experiments involving two or more factors, data were analyzed by one-way, repeated measures ANOVA followed by Dunnett post-hoc multiple comparisons tests. Data were assumed to follow a normal distribution (no tests were performed). Values denoted as follows were considered significant: *, p < 0.05; **, p < 0.01; ***, p < 0.001.

## RESULTS

### Development and Validation of the Method

The Seahorse XF Analyzer provides kinetic readouts of the oxygen consumption rate (OCR; in pmol O_2_/min) and extracellular acidification rate (ECAR; in mpH/min). These measurements very often correlate with oxidative phosphorylation (OCR) and glycolysis (ECAR), but do not directly reflect the quantity of ATP produced via each pathway. A strategy to convert OCR and ECAR readings into rates of ATP produced from oxidative phosphorylation and glycolysis involved three principal steps:

i. Transforming ECAR (mpH/min) into a proton production rate (pmol H^+^/min) so rates of medium acidification could be directly compared against rates of oxygen consumption (pmol O_2_/min) or lactate efflux (pmol lactate/min) with similar units.

a. mpH/min → pmol H^+^/min
ii. Accounting for sources of acidification not associated with glycolysis and fermentation so changes in H^+^ production quantitatively reflect lactate efflux.

a. pmol H^+^/min → pmol H^+^_Lactate_/min
iii. Converting rates of oxygen consumption and lactate efflux into ATP produced from oxidative phosphorylation and glycolysis using established stoichiometry.

a. pmol O_2_/min ® pmol ATP_OxPhos_/min
b. pmol H^+^_lactate_/min ® pmol ATP_Glyco_/min
c. pmol ATP/min = pmol ATP_OxPhos_/min + pmol ATP_Glyco_/min

The work presented uses a 96-well Seahorse XF Analyzer platform. Although some values cannot be directly translated to 24-well instruments due to differences in volume of the measurement microchamber, the experimental strategy to calculate these values is universal across all Seahorse XF platforms.

(i) Transforming ECAR into a proton production rate

It was first necessary to convert ECAR into values of pmol H^+^/min so rates of medium acidification could be easily compared against OCR (in pmol O_2_/min) and parallel lactate efflux assays (in pmol lactate/min). This unit conversion required calculating the buffering power of the experimental medium. Buffering power is defined here as the pH change in the measurement microchamber elicited by a given amount of H^+^ (Fig. 1A). Importantly, sequential additional of H_2_SO_4_ showed the change in pH was directly proportional to the added H^+^ over a range greater than 0.5 pH units (Fig. 1B). Predictably, as the concentration of HEPES in the medium increased, there was a smaller drop in pH elicited from the added H^+^. The results demonstrate that extracellular acidification measurements in mpH/min are readily converted to values of pmol H^+^/min by applying a scalar factor accounting for the buffering power of the medium and the volume of the measurement microchamber. Sample calculations and step-by-step instructions for transforming ECAR into values of pmol H^+^/min are provided in the first tab of the Appendix Worksheet available in the manuscript Appendix.

**Figure 1.**
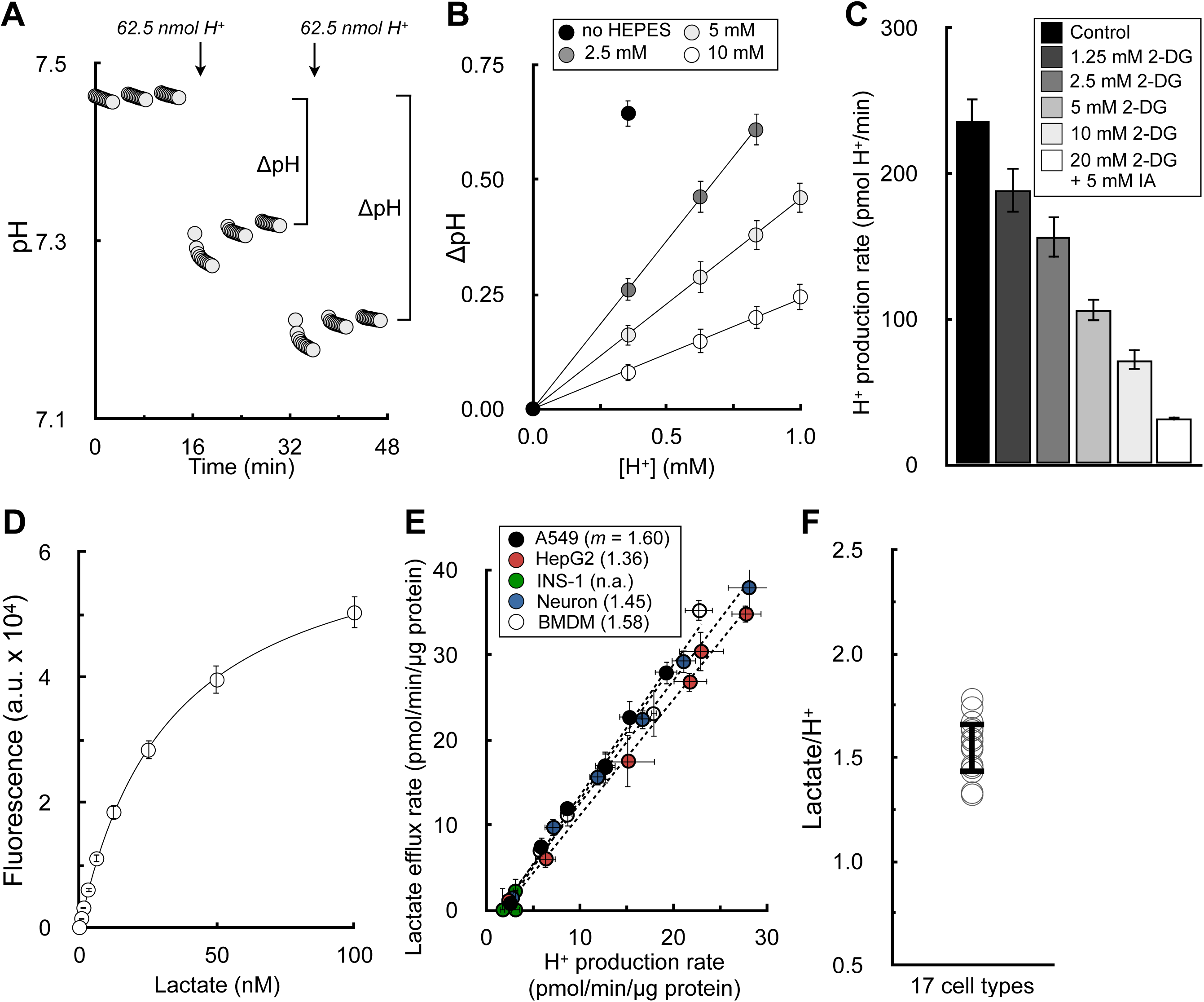
Extracellular acidification is directly proportional to lactate efflux in the absence of respiration. All data are mean ± S.E.M. unless otherwise specified. (A) Sample trace of H_2_SO_4_ injections to determine buffer capacity of Seahorse experimental medium. Error bars are obscured by the symbol. (n=16 technical replicates from a single experiment) (B) Changes in pH from sequential additions of 62.5 nmol H^+^ via the injector ports. The experimental medium is DMEM supplemented with 8 mM glucose, 2 mM glutamine, 2 mM pyruvate, and the indicated concentration of HEPES (pH 7.4). The microchamber volume of the 96-well XF Analyzer is 2.28 μL. Step-by-step calculations are available in the Appendix Worksheet. (n=6 experimental replicates) (C) The H^+^ production rate is given for A549 cells treated acutely (20 minutes prior to assay) with varying concentrations of 2-deoxyglucose (2DG) or 2DG with iodoacetic acid (IAA) to block glycogenolysis. The experimental medium contains 5 mM glucose with 5 mM HEPES, 200 nM rotenone (complex I inhibitor), 1 μM antimycin (complex III inhibitor), and 2 μM oligomycin (complex V inhibitor to block ‘reverse’ hydrolysis of ATP upon respiratory chain inhibition). (n=6 biological replicates) (D) Sample trace of standard curve used to measure lactate collected from experimental medium. (n=3 technical replicates) (E) Plot of lactate efflux rates measured as in (D) against H^+^ production rates measured in the Seahorse XF Analyzer. Experimental conditions and titration of the rate with glycolytic inhibitors are as in (C). Parenthetical values given in the key are the fitted slopes for each individual cell type. (n=6 biological replicates) (F) Aggregate lactate:H^+^ ratios, calculated as in (E) for 17 different cell types, yielding an average value of 1.53 ± 0.12 (mean ± standard deviation).

(ii) Accounting for sources of acidification other than lactate efflux

It was then necessary to determine whether measurements of H^+^ efflux could indeed accurately measure lactate efflux. In principle, there could be several reactions that could cause a meaningful net pH change in the experimental medium in addition to the incomplete oxidation of uncharged glucose into anionic lactate (Fig. EV1A). Other glycolysis-linked reactions can result in medium acidification, such as the efflux of anionic pyruvate or CO_2_ evolution during the oxidative pentose phosphate pathway (Fig. EV1B). Additionally, several mitochondrial dehydrogenases will generate CO_2_ during the complete oxidation of energy substrates and oxidative phosphorylation. These include the 2-oxoacid dehydrogenase family (pyruvate dehydrogenase, α-ketoglutarate dehydrogenase, and the branched chain keto acid dehydrogenase) and isocitrate dehydrogenase (Fig. EV1C). Furthermore, specific cell types may release organic acids under certain condition, such as hepatocyte efflux of ketone bodies during robust fatty acid oxidation or neuronal release of glutamate during prolonged depolarization.

Rather than attempt to account for every possible acidifying reaction, we began by testing the hypothesis that respiratory acidification (CO_2_ evolution from mitochondrial dehydrogenases linked to O_2_ consumption) would be the only acidifying process with a large enough flux to affect the measurements on a scale comparable with lactate efflux in the basal state. If this hypothesis is correct, then the following testable predictions should hold. First, if respiratory acidification is indeed the only measurable source of non-glycolytic acidification, any remaining acid production should be attributable to glycolysis when respiration is inhibited. Similarly, in the absence of glycolysis, respiration should be directly proportional to the remaining acidification rate.

#### In the absence of respiration, glycolysis is the exclusive source of measurable extracellular acidification

We first measured whether lactate efflux was directly proportional to H^+^ production after chemical inhibition of the respiratory chain. In the presence of electron transport chain inhibitors, cellular H^+^ production was gradually reduced with increasing concentrations of glycolytic inhibitors in A549 cells (Fig. 1C). We then measured the lactate content in the spent experimental medium from these samples using an enzymatic lactate assay (Fig 1D) to determine the relationship between lactate efflux and acid production in the absence of respiration (Fig. 1E). As a negative control, no measurable acidification was observed in cultured pancreatic β-cells(Zhao *et al*, 2001), which do not express appreciable levels of lactate dehydrogenase or monocarboxylate transporter-1. Using 17 different cell types comprised of primary and immortalized cultures, this analysis yielded an average lactate:H^+^ ratio of 1.53 ± 0.12 (Fig. 1F; Appendix Table 1).

Unexpectedly, this value deviated substantially from our expectations and was substantially different from previous efforts to correlate H^+^ production and lactate efflux rates (Mookerjee *et al*, 2015). In principle, the lactate:H^+^ ratio should never be greater than 1, since lactate efflux should never exceed the total H^+^ efflux given their 1:1 stoichiometry. In fact, the theoretical lactate:H^+^ should be less than 1 due to acidifying reactions associated with glycolysis but distinct from lactate efflux (Fig. EV1B). We reasoned that this discrepancy could be due to the incomplete coverage of the measurement well by the fluorometric sensor of the Seahorse XF cartridge: although the enzymatic lactate measurements used medium collected from the entire well, outer portions of the well are likely undetected by the fluorometric sensor spot(s) during the acidification measurements (Gerencser *et al*, 2009) (Fig. EV2A).

To test this, we scraped cells from the measurement well from either the center of the three risers directly under the sensor (creating a “donut” shape of cells in the well) or on the rim of the plate outside the three risers (leaving cells in the center untouched) (Fig. EV2B). As predicted, both OCR and ECAR were significantly reduced when cells in the middle of the well were removed, but losing cells along the outer rim did not significantly affect the measurements (Fig. EV2C). To ensure the results were not due to changes in cell number, the lactate:H^+^ ratio was calculated for these conditions. Consistent with the sensor not seeing the outer edges of the well, lactate:H^+^ was lowered close to 1 when cells were removed from the outer rim, and increased beyond 8 when cells were removed from center of the plate (Fig. EV2D). Rather than a systemic error with the buffer capacity or enzymatic measurements, the result demonstrates that incomplete sensor coverage of the measurement well is a main driver of why lactate:H^+^ values deviate from theoretical expectations.

#### In the absence of glycolysis, respiration is the exclusive source of measurable medium acidification

Having shown that medium acidification is tightly correlated with glycolysis in the absence of respiration in Figure 1, we then tested whether medium acidification would be tightly correlated to respiration in the absence of glycolysis. We therefore measured acidification rates in HepG2 cells offered glucose-free medium containing pyruvate, glutamine, and the glycolytic inhibitors 2-deoxyglucose and iodoacetate. The rates of respiration and acidification were titrated by adjusting flux through the electron transport chain (with oligomycin, FCCP, and rotenone with antimycin A) and blocking substrate uptake and oxidation (with varying concentrations of the mitochondrial pyruvate carrier inhibitor UK5099 and the glutaminase inhibitor BPTES) (Figs. 2A&B).

**Figure 2.**
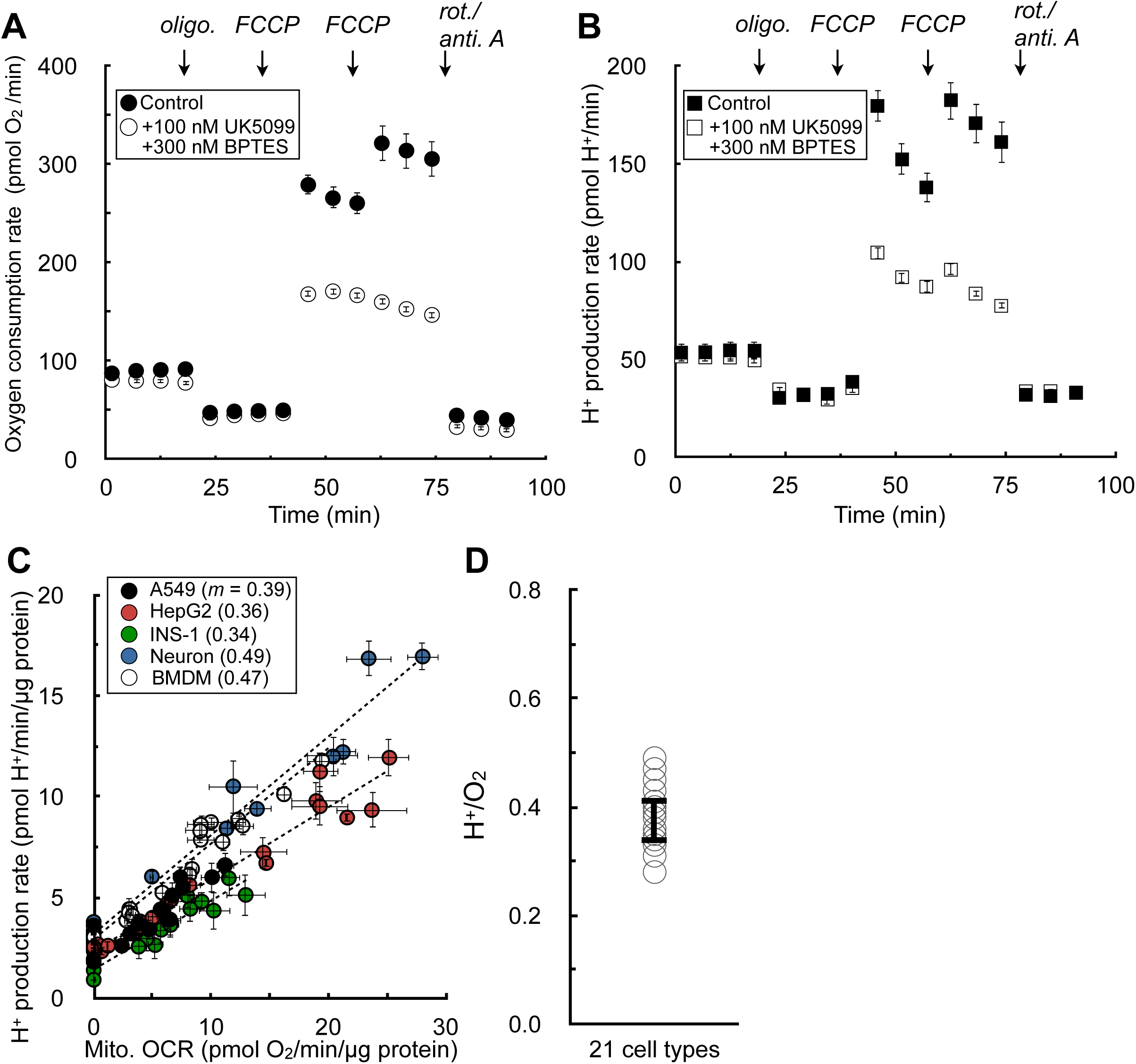
Extracellular acidification is directly proportional to respiration in the absence of glycolysis. All data are mean ± S.E.M. unless otherwise specified. (A) Sample kinetic trace of oxygen consumption in HepG2 cells in medium containing 5 mM pyruvate, 5 mM glutamine, 5 mM HEPES, 2 mM 2-deoxyglucose, and 50 μM iodoacetate. Where indicated, the respiratory rate was modulated with addition of UK5099 (MPC inhibitor to block pyruvate oxidation) and BPTES (glutaminase inhibitor to block glutamine oxidation). (n=8 technical replicates from a single experiment) (B) Sample kinetic trace of H^+^ production rates measured from (A). (n=8 technical replicates from a single experiment) (C) Plot of H^+^ production rates against oxygen consumption rates for experiments conducted as in (A & B). The rates are titrated with mitochondrial effectors (oligomycin, varying concentrations of FCCP, rotenone/antimycin A) or combined treatment with UK5099 and BPTES. Individual points are taken as the average values for all measurements for a particular treatment (e.g. average of three measurements in response to initial FCCP addition). Parenthetical values given in the key are the fitted slopes for each individual cell type. (n=6 biological replicates) (D) Aggregate H^+^:O_2_ ratios, calculated as in (C) for 21 different cell types, yielding an average value of 0.38 ± 0.05 (mean ± standard deviation).

We observed a proportional relationship between respiration and medium acidification in the absence of glycolysis (Fig. 2C). We again extended this analysis to a panel of 21 different primary and immortalized cell preparations to yield an average H^+^:O_2_ ratio of 0.38 ± 0.05 (Fig. 2D; Appendix Table 2). However, similarly to the values obtained for lactate:H^+^, the composite H^+^:O_2_ again deviated sharply from expectations and previous attempts to correct ECAR for respiratory acidification. The value obtained was much lower than the theoretical H^+^:O_2_ of complete oxidation of either pyruvate (0.80) or glutamine (0.67) (Mookerjee *et al*, 2015).

As such, we conducted a series of experiments to better understand the factors that shape the H^+^:O_2_ value and whether experimental perturbations would change respiratory acidification in expected ways. We first examined whether oxidation of different substrates would perturb H^+^:O_2_ values, as complete oxidation of various substrates yields varying H^+^:O_2_ ratios. Indeed, the H^+^:O_2_ ratio resulting from pure glutamine oxidation was fractionally lower than pure pyruvate oxidation (Figs. EV3A-C).

We then further examined whether choice of respiratory substrates could influence H^+^:O_2_ ratios in isolated mitochondria, a simplified system where the experimental conditions are highly controlled and metabolic reactions can be well defined. We tested acidification from uncoupled respiration in rat heart mitochondria offered (i) pyruvate with malate (P/M), (ii) P/M and inhibitors of multiple enzymes creating a truncated TCA cycle (P/M truncated) (Mookerjee *et al*, 2015), and (iii) succinate with rotenone (S/R). The results qualitatively exhibited the expected behavior: the H^+^/O_2_ ratio for mitochondria oxidizing P/M increased in the presence of inhibitors which blocked downstream TCA cycle reactions (e.g. the alkalinizing succinyl CoA synthetase reaction). Moreover, succinate oxidation in the presence of rotenone, which will support oxygen consumption without generating CO_2_, shows robust respiratory rates with negligible changes in pH (Figs. EV3D,E).

However, even in this reductionist system, the measured H^+^:O_2_ fell far below expected values (theoretical H^+^:O_2_ of P/M truncated = 1). We therefore used the XF24 system to further ensure that the result was not a peculiarity of an individual instrument platform. Using the same workflow as before, and accounting for the larger microchamber volume of the 24-well platform, nearly identical results were observed where respiratory acidification in isolated mitochondria fell short of theoretical predictions (Fig. EV3F).

Lastly, in addition to showing that respiratory substrates could affect the H^+^:O_2_ ratio, we wanted to determine whether forced efflux of organic acids other than lactate could also affect the measurements in predictable ways. For a proof-of-concept example, we measured whether NMDA-induced efflux of glutamate would alter the H^+^:O_2_ in primary cortical neurons. Neurons were offered exclusively β-hydroxybutyrate with glycolytic inhibitors, so lactate efflux or sources of glycolytic acid production could not interfere with the measurements. As expected, NMDA-induced efflux of glutamate in primary cortical neurons caused a pronounced spike in the acidification rate (Fig. EV3G) and the H^+^:O_2_ ratio (Fig. EV3H) that was sensitive to the NMDA-receptor antagonist MK801(Divakaruni *et al*, 2017). The result was not attributable to glycolysis, as lactate efflux was negligible and unchanged across conditions (Fig. EV3I). Taken together, the demonstration that H^+^/O_2_ ratios respond in expected ways to changes in substrate oxidation and non-glycolytic acid efflux provided additional confidence in the accuracy of the measured values.

(iii) Converting rates of oxygen consumption and lactate efflux into ATP produced from oxidative phosphorylation and glycolysis

The results from Figures 1 and 2 demonstrate that rates of H^+^ production can, in principle, quantify lactate efflux after correcting for respiratory acidification and accounting for the sensor coverage of the plate well. We therefore transformed H^+^ production rates into lactate efflux rates in Equation 1 as follows:

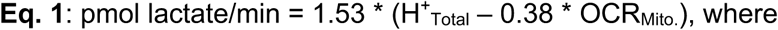

- 1.53 = average lactate:H^+^ value obtained in Fig. 1;
- H^+^_Total_ = H^+^ production rates read by the instrument;
- 0.38 = average H^+^:O_2_ value obtained in Fig. 2;
- OCR_Mito._ = basal mitochondrial respiration;

To validate whether these average values obtained under constrained conditions could be broadly applied across different cell types under typical assay conditions (i.e. complete assay medium with both glycolysis and oxidation phosphorylation functional), we compared lactate efflux calculated using the XF Analyzer against an enzymatic assay in A549 adenocarcinoma cells, C2C12 myoblasts, and murine BMDMs (Fig. 3A). Prior to adjusting the measurements as in Eq. 1, the H^+^ production rate underestimated the rate of lactate efflux (comparing leftmost bars in each graph with rightmost). But as expected, correcting Seahorse XF measurements for respiratory acidification and sensor coverage resulted in lactate efflux values (middle bars) that matched enzymatic measurements. Taken together with Figure 1, the results suggest the consensus values and approximations used here should be applicable across a broad range of cell types under normal assay conditions.

**Figure 3.**
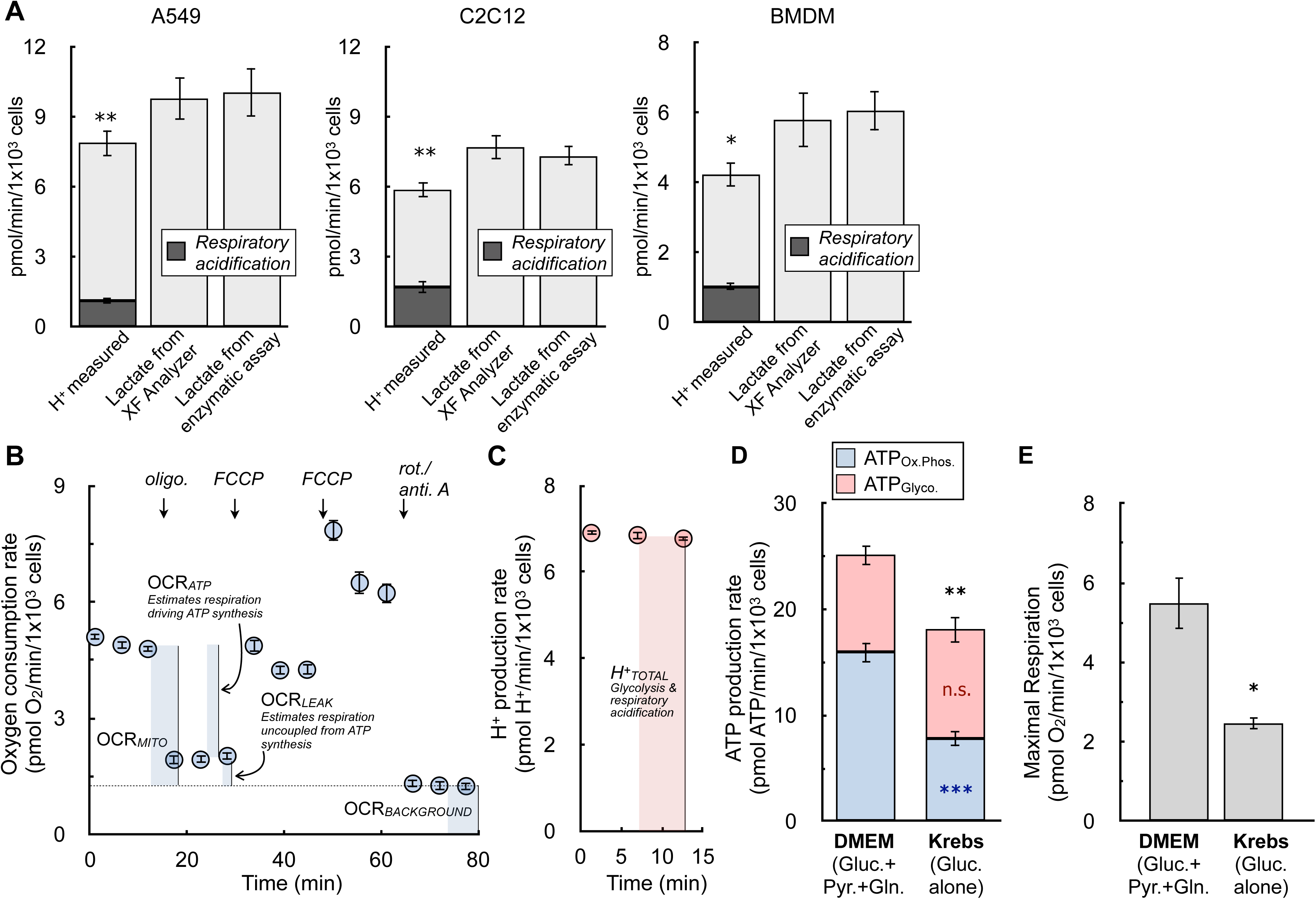
Transformation of ECAR into lactate efflux allows calculation of ATP production rates. All data are mean ± S.E.M. unless otherwise specified. (A) The H^+^ production rate from the XF Analyzer (left bar in each graph), the calculated lactate efflux rate from the XF Analyzer (center bar), and the lactate efflux rate measured from an enzymatic assay (right bar) are given for A549 cells, C2C12 cells, and primary bone marrow-derived macrophages (BMDMs). Cells are offered 8 mM glucose, 2 mM pyruvate, and 2mM glutamine in experimental medium supplemented with 5 mM HEPES. The shaded component of the H^+^ production rates for each cell type is the calculated contribution from respiratory acidification. (n≥4 biological replicates) (B) The oxygen consumption rate from a representative experiment with A549 cells is presented. Assay medium is as in (A). The parameters needed to calculate the ATP production rate are indicated with in the figure. (n=6 technical replicates) (C) The H^+^ production rate from the kinetic trace in (B) is presented. Only the initial rates are shown, as these are the only measurements required to calculate the ATP production rate. (n=6 technical replicates) (D) ATP production rates calculated for A549 cells in DMEM supplemented with 8 mM glucose, 2 mM glutamine, and 2 mM pyruvate and Krebs-Henseleit buffer supplemented with only 10 mM glucose. (n=4 biological replicates) (E) Maximal respiratory rates measured in response to oligomycin and FCCP for treatments as in (D). (n=4 biological replicates)

The ability to quantify lactate efflux with rates of H^+^ production allows the transformation of ECAR into a rate of ATP produced by glycolysis. All necessary parameters are obtained during standard respirometry profiling (Figs. 3B & C), and sample calculations with representative data sets are available in the Appendix Worksheet.

The rate of mitochondrial ATP production can be estimated by applying a P:O ratio (moles of ADP phosphorylated per mole of oxygen consumed) to the respiration that drives ATP synthesis(Brand, 2005), which is determined using sensitivity to the ATP synthase inhibitor oligomycin. As detailed elsewhere, oligomycin-sensitive respiration will slightly underestimate ATP-linked respiration (due to overestimating proton leak), though this is a small, ∼10% overestimate which can be adjusted (0.1*OCR_Leak_ in Eq. 2) (Affourtit & Brand, 2009). Moreover, assumptions must be made for the selection of P/O ratio based on which substrate is being oxidized. Here, we use a value of P:O = 2.73 (P:O_2_ = 5.45), the maximum value for complex I-driven respiration(Watt *et al*, 2010) and within the range of values for complete glucose oxidation(Mookerjee *et al*, 2017). The rate of ATP from oxidative phosphorylation can therefore be estimated as in Eq. 2:

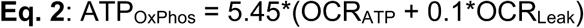

- 5.45 = P:O_2_ (moles of ADP phosphorylated per O_2_ consumed);
- OCR_ATP_ = ATP-linked respiration (respiration sensitive to oligomycin);
- OCR_Leak_ = Respiration associated with proton leak (respiration insensitive to oligomycin);

The rate of glycolytic ATP production can be estimated with 1:1 stoichiometry to the lactate efflux rate (**Eq. 1**), and the total ATP production rate is given by the sum of Eqs. 1 & 2. See Appendix Worksheet for sample calculations

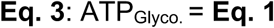

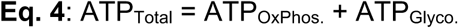

Notably, this calculation will slightly underestimate rates of glycolytic ATP, as it fails to account for the ATP generated during glycolysis when glucose-derived pyruvate is funneled into mitochondria for complete oxidation (Mookerjee *et al*, 2017). Accounting for this would require knowing (or assuming) the fractional contribution of glucose oxidation to oxidative phosphorylation relative to other substrates. To test whether this omission would appreciably affect experimental conclusions, we measured rates of ATP production in A549 cells offered complete medium or a simple salts medium supplemented with only glucose. Offering cells only glucose in simple salts allowed calculation of this additional glycolytic ATP by applying known stoichiometry and assuming respiration is exclusively driven by glucose oxidation with no contribution from endogenous substrate stores. However, rates of both oxidative phosphorylation and uncoupler-stimulated respiration were substantially limited when cells were offered medium containing solely glucose (Figs. 3D&E). Additionally, rates of glycolytic ATP production between groups were only marginally altered despite accounting for the cytoplasmic ATP production linked to mitochondrial glucose oxidation (Fig. 3D). We therefore chose not to account for this component, favoring – on balance – an approach where cells are offered complete DMEM to measure rates of oxidative phosphorylation and uncoupled respiration that are not restricted by substrate limitation.

Lastly, prior to applying the method to various models of cell activation, we measured how altered rates of organic acid efflux in response to activation stimuli may affect the measurements. We chose adrenergic activation of brown adipocytes as a model system, as adipocyte lipolysis is known to cause release of fatty acids into the experimental medium (Thompson *et al*, 2010). Figs. EV4A-C show that stimulation of brown adipocytes with norepinephrine results in the hallmark increase in uncoupled respiration, as well as an increased rate of H^+^ production. To determine whether the contribution of organic acid efflux was different upon norepinephrine stimulation, lactate efflux was measured enzymatically from matched samples on the same Seahorse XF plate. As before, comparing lactate efflux calculated from the XF Analyzer against that measured from an enzymatic assay was within experimental error of multiple biological replicates (Fig. EV4D). However, it was clear that non-glycolytic acid production was increased upon norepinephrine stimulation: the XF Analyzer showed almost a two-fold increase in lactate production in response to adrenergic activation, while the enzymatic assay measured only a 1.5-fold increase (Fig. EV4E). This suggests that H^+^ measurements were somewhat confounded by efflux of organic acids distinct from lactate.

### Application of the Method

#### ATP production rates show patient-derived glioma xenografts acquire an artificial glycolytic phenotype during in vitro cell culture

A clear benefit of measuring ATP production rates is the ability to extend beyond commonly used OCR:ECAR ratios to directly quantify changes in the balance between oxidative phosphorylation and glycolysis. To illustrate this, we analyzed tumor cells from patient-derived glioma xenografts either immediately harvested and purified from mice (∼3 hr. isolation) or matched cells propagated in *in vitro* gliomaspheres (>1 week culture period). *In vivo* evidence suggests that the enhanced aerobic glycolysis observed in some glioma cell culture models may be driven, in part, by artifacts of cell selection or *in vitro* growth conditions (Marin-Valencia *et al*, 2012). To explore this, we again confirmed that our measurements of lactate using H^+^ efflux matched enzymatic measurements for these patient-derived cultures (Fig. 4A). We then examined the information that can be gained from ATP production rates as opposed to OCR:ECAR ratios. In five different cell lines, *in vitro* cultured glioma cells showed a significantly reduced OCR:ECAR ratio compared to matched cells assayed immediately after purification from corresponding murine xenografts (*ex vivo*), indicating a relative shift towards glycolysis in *in vitro* cultured cells (Figs. 4B).

**Figure 4.**
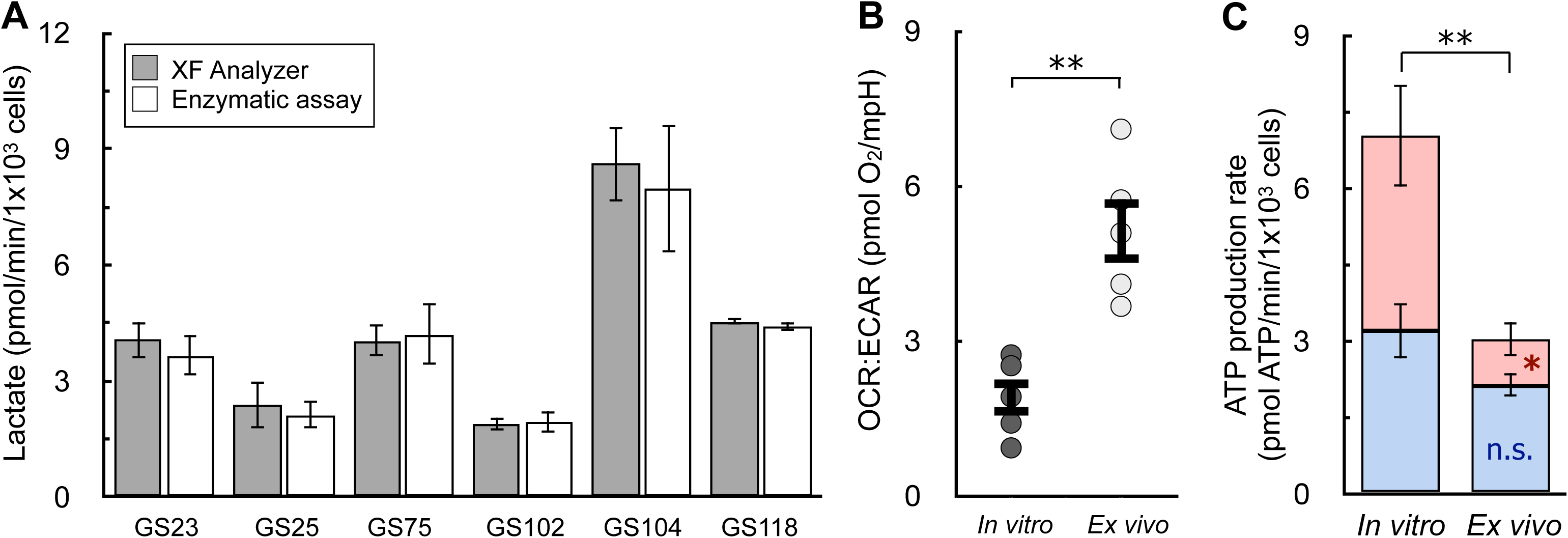
ATP production rates capture changes OCR:ECAR ratios cannot in primary cancer cell models. All data are mean ± S.E.M. unless otherwise specified. *, p<0.05; **, p<0.01. (A) Lactate as measured by the XF Analyzer matches enzymatic lactate measurements for a panel of patient-derived, gliomasphere (GS) cell lines after *in vitro* culture. (n≥3 biological replicates for each line) (B) OCR:ECAR ratios for patient-derived glioblastoma cells cultured *in vitro* or assayed immediately after harvesting from mouse xenographs (*ex vivo*). Cell lined used were GS25, GBM39, GS114, GS116, GS122. (C) ATP Production rates from cell types as in (B).

Quantifying ATP production rates, however, extends this qualitative analysis. Glioma cells analyzed *ex vivo* just after isolation and purification are highly dependent on oxidative phosphorylation (∼70% of total ATP). When cultured *in vitro*, however, cells undergo a remarkable shift in metabolism, increasing their energy demand and switching to glycolysis as the dominant ATP-generating pathway (Figs. 4C). Importantly, there is only a slight, statistically insignificant trend towards increased oxidative phosphorylation, suggesting a specific reprogramming of glycolysis acquired upon *in vitro* culture and not simply an increased energy demand met by both pathways. The result shows that calculating ATP production from oxidative phosphorylation and glycolysis yields insights absent in qualitative, ratio-based analyses, and further supports an *in vivo* bioenergetic role of mitochondria in glioblastoma (Maher *et al*, 2012; Mashimo *et al*, 2014).

#### ATP production rates demonstrate oxidative phosphorylation and glycolysis are differentially engaged upon pro-inflammatory macrophage activation

We then applied our calculation to study pro-inflammatory macrophage activation, another form of immune cell activation associated with substantial metabolic alterations. Upon classical, “type I” activation [LPS (± interferon-γ)], murine BMDMs profoundly remodel energy metabolism away from oxidative phosphorylation (Van den Bossche *et al*, 2016) (Fig. 5A). ATP production is shifted to glycolysis and mitochondria are repurposed to generate signals that support the pro-inflammatory response (Mills *et al*, 2016; West *et al*, 2011; Lampropoulou *et al*, 2016). By measuring this switch with a qualitative OCR:ECAR ratio, metabolic reprogramming with LPS was indistinguishable from respiratory chain inhibition with complex I inhibitor rotenone (Fig. 5B, inset). However, quantifying ATP production rates per cell revealed an increased energy demand associated with LPS activation, discriminating between mitochondrial dysfunction and mitochondrial repurposing during macrophage activation (Fig. 5B; Fig. EV5).

**Figure 5.**
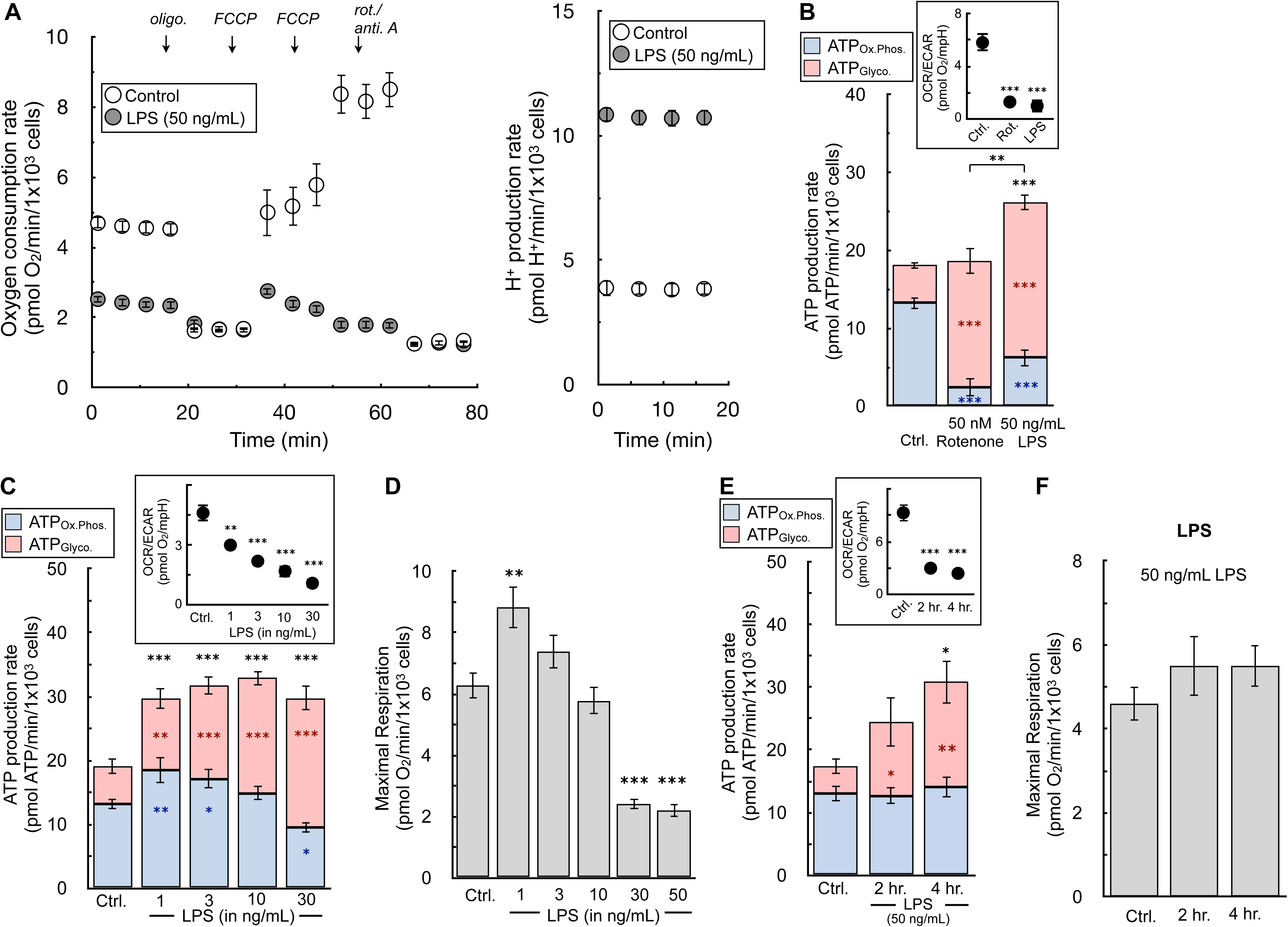
ATP production rates highlight independent regulation of glycolysis and oxidative phosphorylation during LPS activation of macrophages. All data are mean ± S.E.M. unless otherwise specified. *, p<0.05; **, p<0.01; ***, p<0.001. (A) The oxygen consumption rate and H^+^ production rate from a representative experiment with BMDMs in response to treatment with 50 ng/mL LPS for 24 hr. (n=5 technical replicates) (B) The ATP production rate is calculated for BMDMs in response to 50 ng/mL LPS or 50 nM rotenone for 24 hr. (*Inset*) The OCR:ECAR ratio is presented for each condition. (n=6 biological replicates) Representative images used for cell counts are provided in the supplementary material. (C) The ATP production rate is calculated for BMDMs in response to varying concentrations of LPS ranging from 1-30 ng/mL for 24 hr. (*Inset*) The OCR:ECAR ratio is presented for each condition. (n=5 biological replicates) (D) FCCP-stimulated, maximal respiratory rates from treatments as in (C). (n=5 biological replicates) (E) The ATP production rate is calculated for BMDMs in response to treatment for 2 or 4 hr. with 50 ng/mL LPS. (*Inset*) The OCR:ECAR ratio is presented for each condition. (n=4 biological replicates) (F) FCCP-stimulated, maximal respiratory rates from treatments as in (E). (n=4 biological replicates)

Quantifying ATP production rates over a range of LPS concentrations further highlights the depth of information gained from the measurements. Qualitative OCR:ECAR ratios in response to increasing LPS concentrations (1-30 ng/mL) suggest a uniform, coordinated shift away from oxidative phosphorylation towards glycolysis to meet ATP demands upon activation (Fig. 5C, inset). Cellular ATP production rates, on the other hand, surprisingly reveal these two pathways are engaged independently. Increases in glycolysis and overall ATP utilization are observed with treatments as low as 1 ng/mL LPS, whereas rates of mitochondrial ATP production are not compromised until LPS concentrations rise above 10 ng/mL (Fig. 5C). Additionally, rates of maximal respiration are not compromised at low concentrations of LPS that substantially increase glycolysis, further demonstrating that the increase in glycolytic ATP is not simply a response to impaired mitochondrial oxidative metabolism (Fig. 5D).

Indeed, the time-dependence of energetic activation further reveals that regulation of mitochondrial and glycolytic ATP production are disengaged: glycolysis increases within two hours of activation while rates of oxidative phosphorylation and maximal respiration are maintained up to at least four hours (Figs. 5E&F). The results show the hallmark metabolic changes upon pro-inflammatory macrophage activation are a composite phenotype where increases in ATP demand and glycolysis are controlled distinctly from reductions in oxidative phosphorylation.

#### ATP production rates can measure healthy bioenergetic changes during cell activation that are lost with static ATP measurements

We finally applied this analysis to models of acute cell activation to highlight the differences between ATP production rates and measurements of ATP content. Healthy cell activation such as neuronal plasma membrane depolarization or T cell receptor activation should minimally affect snapshot-in-time intracellular ATP levels, as cells will readily adapt to a new steady-state of increased ATP turnover (Brand & Nicholls, 2011). In primary cortical neurons, for example, addition of the Na^+^- channel activator veratridine stimulates Na^+^/K^+^-ATPase activity (Fig. 6A, top). Both oxygen consumption and extracellular acidification rates increased in response to acute veratridine addition (Choi *et al*, 2009) (Fig. 6A, bottom). Quantifying ATP production rates showed almost a three-fold increase in the overall rate of ATP turnover upon depolarization, with the balance between oxidative phosphorylation and glycolysis broadly unchanged upon activation (Fig. 6B). To demonstrate the sensitivity of the assay, we used a sub-saturating amount of the Na^+^/K^+^-ATPase inhibitor ouabain (2 nM) to demonstrate the assay can pick up small but meaningful changes in the cellular ATP demand. Substantial change in steady-state ATP utilization, however, did not appreciably change static ATP levels, though levels were predictably altered in response to metabolic inhibitors coupled with neuronal depolarization (Fig. 6C).

**Figure 6.**
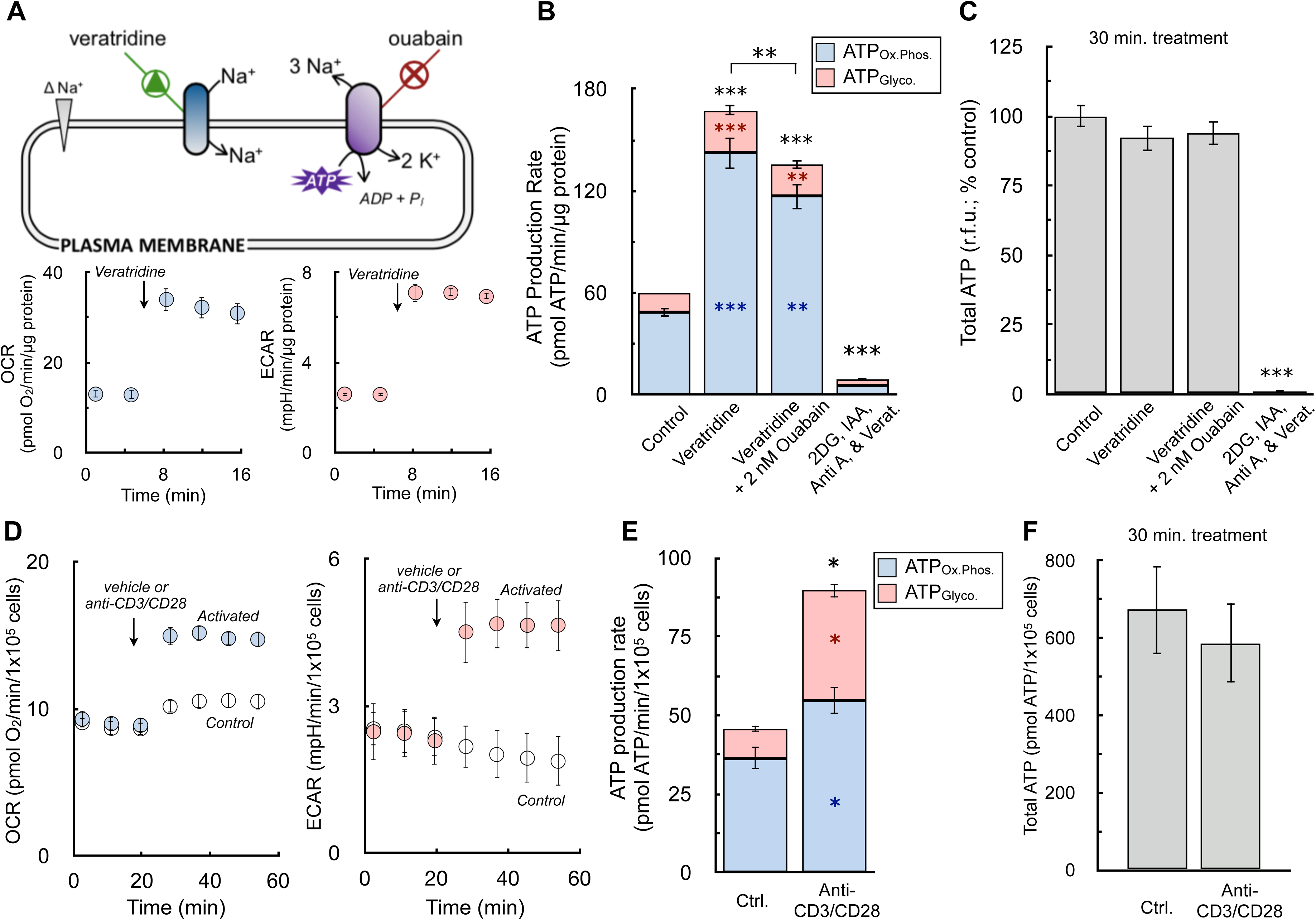
ATP production rates capture healthy changes in cell activation that static ATP measurements cannot. All data are mean ± S.E.M. unless otherwise specified. *, p<0.05; **, p<0.01; ***, p<0.001. (A) (*Top*) Schematic depicting increase in Na^+^/K^+^-ATPase activity by veratridine, which will depolarize the plasma membrane by increasing sodium influx, thereby triggering ouabain-sensitive increases in ATP production. (*Bottom*) Sample kinetic traces of OCR and ECAR in primary neurons upon acute addition of 1.5 μM veratridine. (n=10 technical replicates) (B) ATP production rates from oxidative phosphorylation (ATP_Ox.Phos._, blue) and glycolysis (ATP_Glyco._, red) in response to (*left to right*) vehicle, 1.5 μM veratridine, veratridine plus 2 nM ouabain to demonstrate sensitivity of the assay, and veratridine plus 1 μM antimycin, 5 mM 2-deoxyglucose, and 1 mM iodoacetic acid to cause an energy crisis and deplete ATP levels. (n=4 biological replicates) (C) Total ATP levels measured by luminescence after 15 min treatments as in (B). (n=4 biological replicates) (D) Sample kinetic traces of OCR and ECAR in murine T cells acutely activated with anti-CD3/CD28. (n=4 technical replicates) (E) ATP production rates from oxidative phosphorylation and glycolysis in response to anti-CD3/CD28. (n=3 biological replicates) (F) Total ATP levels measured by luminescence after 30 min activation as in (E). (n=3 biological replicates)

To further demonstrate the principle that ATP production rates provide information during cell activation lost with steady-state measurements, we measured human CD4^+^ T cell activation in response to anti-CD3/CD28 stimulation. Here again, acute cell activation increased rates of both OCR and ECAR (Fig. 6D). As is well documented, T cell activation caused a pronounced increase in glycolysis to meet the energetic and biosynthetic demands of proliferation, cytokine production, and clonal expansion (Maciver *et al*, 2013; Buck *et al*, 2017). Nonetheless, despite a four-fold increase in glycolytic ATP production, comparing both pathways showed oxidative phosphorylation remained the primary source of cellular ATP production, reinforcing an important role for mitochondria during early T cell activation (Sena *et al*, 2013) (Fig. 6E). As was demonstrated previously for neurons, a physiological increase in the ATP demand that is readily met by T cells did not manifest in substantially altered ATP levels (Fig. 6F). In total, the results show real-time ATP production rates can reveal activation-associated changes in energy metabolism that static ATP levels cannot.

## DISCUSSION

### ATP fluxes during cell activation

Our empirical approach to adjust proton efflux rates for respiratory acidification enables quantification of lactate efflux using OCR and ECAR. Furthermore, obtaining quantitative lactate readouts from the XF Analyzer allows estimation of cellular ATP production rates. ATP production rates from both oxidative phosphorylation and glycolysis provide additional insight into how ATP utilization is distributed during cell adaptation and activation. For example, *in vivo* stable isotope tracing has shown that *in vitro* models of glioblastoma and other cancers can overestimate the contribution of glycolysis to energy metabolism (Faubert *et al*, 2017; Marin-Valencia *et al*, 2012; Momcilovic *et al*, 2019). In support of this, matched samples of patient-derived glioblastoma cells clearly show the acquisition of an artificially high ATP demand driven almost entirely by a five-fold increase in glycolysis in cells cultured *in vitro* relative to those assayed immediately after isolation from murine orthotopic xenografts.

Additionally, the assay reveals that cells can adjust the balance of ATP production differently depending on the type of activation and biological context. As an example, acute activation of the Na^+^/K^+^-ATPase in primary cortical neurons substantially increases the ATP demand in a way that does not broadly change the proportion of ATP produced by oxidative phosphorylation and glycolysis. Despite an acute, 2.5-fold increase in ATP production, the balance between the two pathways remained roughly constant with about 85% of the total ATP produced by mitochondria. It is sometimes stated that the Na^+^/K^+^-ATPase and other plasma membrane ATPases preferentially rely on ATP produced by glycolysis (Epstein *et al*, 2014; Sepp *et al*, 2014). However, our results do not support this relationship, at least in two-dimensional cell culture models not subject to microenvironmental gradients.

Unlike the instantaneous increase in ATP demand elicited by plasma membrane depolarization, immune cell activation shows that cells can redistribute the balance between oxidative phosphorylation and glycolysis when activating appropriate signaling cascades. Both T cell antigen receptor (TCR)-linked activation in human T cells as well as TLR-mediated activation of murine macrophages demonstrated that the increase in ATP utilization upon cell activation was increasingly met by glycolysis relative to the naïve state. In T cells this increased glycolytic flux supports generation of biosynthetic intermediates necessary for cell growth and expansion (Maciver *et al*, 2013). Given that many types of TLR activation cause growth arrest in murine macrophages, the observed increase in glycolysis serves a different role. For example, the increase in glycolysis is likely associated with increased flux through the pentose phosphate pathways to support cytoplasmic NADPH production and fuel the anti-microbial NADPH oxidase (Cathcart, 2004). In fact, the approach highlights that regulation of glycolysis and oxidative phosphorylation are disengaged during the pro-inflammatory activation of macrophages. Rather than a coordinated process where mitochondrial ATP production decreases alongside a matched increase in glycolysis, pro-inflammatory stimuli can substantially increase glycolysis without changing rates of oxidative phosphorylation.

### Adjusting proton production rates to quantify lactate efflux

In line with previous reports, our empirical approach determined that glycolytic and respiratory acidification were the only dominant contributors to the extracellular acidification rate under normal assay conditions. By and large, other reactions that could result in a net pH change of the medium and alter the H^+^ efflux rate either (i) did not have with fluxes comparable to those of respiratory and glycolytic acidification, or (ii) occurred proportionally with respiratory CO_2_ evolution or lactate efflux (e.g. CO_2_ evolution from the pentose phosphate pathway) and fell within a reasonable margin of experimental error. Cell types or experimental conditions that may deviate from this framework are addressed in the “Best practices” section of the Discussion.

Although we demonstrate that the H^+^ efflux rate can quantitatively measure lactate efflux after accounting for respiratory CO_2_, the empirical values obtained did not follow the expected stoichiometry for either lactate:H^+^ or H^+^:O_2_ and were inconsistent with prior work (Mookerjee *et al*, 2015). These deviations from expected stoichiometry presented here, however, have theoretical and experimental support. The value of 1.53 ± 0.12 for the lactate:H^+^ can be explained in part by incomplete coverage of the plate well by the fluorometric sensor in the XF cartridge, as cells in the center of well have a far greater effect on the readings than those on the rim.

In fact, the expected lactate:H^+^ value of ∼1 is obtained when cells are only within the circle connecting the three microplate risers, roughly corresponding to the XF96 fluorometric sensor area. The demonstration that cells directly under the measurement sensor are preferentially, if not exclusively, read by the cartridge suggests that there is no systematic error with the enzymatic lactate measurements. Additionally, the result implies a 1:1 lactate:H^+^ ratio should never have been expected, particularly in older instruments where the fluorometric sensors consist of two small spots (Gerencser *et al*, 2009) and not the larger smears found in newer instruments. Seahorse XF Wave software now incorporates this correction into the Proton Efflux Rate (PER) parameter for all multi-well platforms, accounting for the buffering power of the medium and incorporating the geometric corrections obtained from this work.

The H^+^:O_2_ values from our analysis also deviate from theoretical stoichiometries expected from the complete oxidation of known substrates. Nonetheless, the obtained values change with the appropriate directionality and magnitude in response to oxidizing varying energy substrates in both intact cells and isolated mitochondria. Moreover, lactate efflux rates calculated from the XF Analyzer using an H^+^:O_2_ of 0.38 ± 0.05 to adjust for respiratory CO_2_ matched enzymatic lactate assays, suggesting there was no systemic error in measuring buffering power of the experimental medium.

Notably, the discrepancy between the obtained and expected H^+^:O_2_ values persists in isolated mitochondria, where the experiments are highly controlled and the metabolic reactions are well defined. In this system, the apparent H^+^:O_2_ values for a truncated reaction where mitochondria are offered pyruvate and malate to generate citrate (theoretical H^+^:O_2_ = 1 (Mookerjee *et al*, 2015)) are more than 3-fold lower than what is expected in principle and previously observed. The result is likely due, in part, to the different ways the XF Wave software calculates rates of H^+^ efflux and O_2_ consumption. While the reported rates of H^+^ production are a straightforward measurement, the instrument reports rates of O_2_ consumption only after an empirical correction to deconvolute the effects of back-diffusion of ambient oxygen into the measurement microchamber(Gerencser *et al*, 2009). This includes use of an ‘apparent chamber volume’ that is 3-times greater than the actual measurement chamber volume to account for oxygen storage in the walls of the polystyrene microplate, and the algorithmic constants were determined by matching respiration rates obtained from synaptoneurosomes in the XF Analyzer with values obtained from a platinum-based oxygen electrode (Gerencser *et al*, 2009).

Crucially, these corrections would necessarily account, at least partially, for respiration in material along the rim of the plate unseen by the measurement sensor because they are clamped to values obtained from a Clark-type electrode. However, no such corrections for material unseen by the measurement sensor are made for rates of medium acidification. As such, it is expected that rates of H^+^ production are consistently underestimated relative to rates of O_2_ consumption, and that the observed H^+^:O_2_ ratios fall below theoretical predictions. Put simply, while software corrections adjust OCR to reflect the full well, this is not the case when measuring H^+^ production.

### Assumptions and limitations in calculating ATP production rates

Notwithstanding the deviation from expected values, the consensus values obtained for lactate:H^+^ and the H^+^:O_2_ accurately adjust for respiratory CO_2_ and can be used to quantitatively reflect lactate efflux. They can therefore be used to convert oxygen consumption rates and H^+^ efflux rates into estimates of ATP production as detailed in the Appendix Worksheet.

Any approach to estimate ATP production rate, be it the empirical approach presented here or ones based on theoretical principles (Mookerjee *et al*, 2017), are subject to limitations and rely on multiple assumptions. Our empirical approach uses an average H^+^:O_2_ value for use with standardized experiment medium for cellular bioenergetics: DMEM supplemented with glucose, glutamine, and pyruvate. This approach, however, precludes the ability to estimate ATP supply changes upon shifts in cellular substrate preference (e.g. from fatty acids to glucose) in response to external cues. In principle, this could be done by strictly defining the medium composition and applying different H^+^:O_2_ values based on assumptions or measurements of what substrates are being oxidized.

Additional limitations associated with our approach are (i) use of an approximate P:O value and (ii) an underestimation of glycolytic ATP as described in the results. Both result from conducting experiments in a composite medium more like culture conditions with multiple oxidizable substrates available. It is therefore difficult to know, and perhaps unwise to assume, the fractional contribution of individual substrates to the oxygen consumption rate.

We therefore chose to use an approximate P:O ratio and not to account for glycolytic ATP produced during the complete oxidation of glucose. This approach did not significantly alter measurements of glycolytic ATP in A549 lung adenocarcinoma cells, but the experimental conditions substantially increased rates of oxidative phosphorylation and respiratory capacity. The pre-set calculations of the Seahorse Analytical Software Tools follow this framework: they incorporate average H^+^:O_2_ and lactate:H^+^ values, use an approximate P:O ratio, and ignore the glycolytic ATP contribution from complete glucose oxidation.

### Best practices for calculating of ATP production rates

Many of the best practices associated with this particular assay are shared with Seahorse XF assays in general. These include optimizing the plating density for suitable rates, ensuring all measurements are conducted with rates at a relatively stable steady-states, using the inner 60 wells of the XF96 microplate to avoid temperature and evaporative effects, and following best practices for normalization (Divakaruni & Jastroch, 2022). Of course, any acute compound treatments should precede the injection of rotenone and antimycin A.

When calculating the buffering power of the experimental medium, it is always best practice to use a non-volatile acid (e.g. H_2_SO_4_) or two distinct calibrations (e.g. one with HCl and one with H_2_SO_4_) to minimize instrument and operational error. The buffering power should be checked or recalculated when critical parts of the measurement process are changed – such as switching cartridge lot numbers, batches of medium, or instruments – but does not necessarily need to be calculated alongside each experimental run. In the 96-well instrument, it is often convenient to use columns 1 and 12 of the microplate for these calculations as is shown in the Appendix Worksheet.

Additionally, it may be that the approximations used for this method are substantively inaccurate in specific cases depending on the experimental hypothesis and quantitative rigor required. For example, rates of lactate efflux may be particularly low when using acutely isolated primary cells, and therefore the non-zero background rate of H^+^ efflux (Figs. 1E and 2C) may represent a substantial component of the signal and skew results. Furthermore, conditions where the rate of lactate efflux is matched by organic acid efflux of the same magnitude may also be less amenable to using consensus values of lactate:H^+^ and H^+^:O_2_. Indeed, this is apparent in noradrenaline-stimulated adipocytes known to release fatty acids: our XF Analyzer calculations based on H^+^ release estimated an almost two-fold increase in lactate efflux, whereas the enzymatic assay showed a 1.5-fold increase (Fig. EV4). Additionally, under extreme, non-physiological conditions such as neurons depolarized in the absence of glucose, release of glutamate and other neurotransmitters resulted in a profound increase in acidification entirely independent of lactate efflux.

As such, any system in which the investigator believes that non-lactate acid efflux could prohibitively alter the conclusions should consider independently calculating lactate efflux with other methods such as enzymology or mass spectrometry. However, the empirical approach presented here provides the framework for researchers to calculate lactate:H^+^ and H^+^:O_2_ values tailored to any (monolayer) model system or experimental conditions.

## Supporting information

Appendix Worksheet

## ACKNOWLEDGEMENTS

A.S.D. is supported by National Institutes of Health (NIH) Grants R35GM138003, P30DK063491, and P50CA092131, as well as the W.M. Keck Foundation. B.R.D. is supported by the U.S. Department of Defense National Defense Science and Engineering (NDSEG) Fellowship Program. A.E.J. was supported by the UCLA Tumor Cell Biology Training Program (T32 CA009056) and a Eugene V. Cota Robles Scholarship. A.B.B. was supported by the UCLA Chemistry-Biology Interface Training Grant (T32GM136614). S.J.B. is supported by NIH Grants P01HL146358 and R01HL157710.

## AUTHOR CONTRIBUTIONS

A.S.D. and N.R. supervised all aspects of the project. A.S.D., N.R., S.J.B., A.N.M., B.P.D., D.A.N., A.N., D.A.F., C.A. and O.S. conceptualized the study and/or designed the research plan. A.S.D., B.R.D., K.K.O., A.B.B., W.Y.H., A.E.J., P.S., D.M., A.J.B., L.S., G.W.R., and N.R. conducted experiments. A.S.D., B.R.D., K.K.O., A.B.B., W.Y.H., A.E.J., P.S., and N.R. analyzed the data. A.S.D., B.R.D., and N.R. prepared the figures and wrote the manuscript with editing and approval from all authors.

## MANUSCRIPT EXPANDED VIEW

**Expanded View Figure 1.**
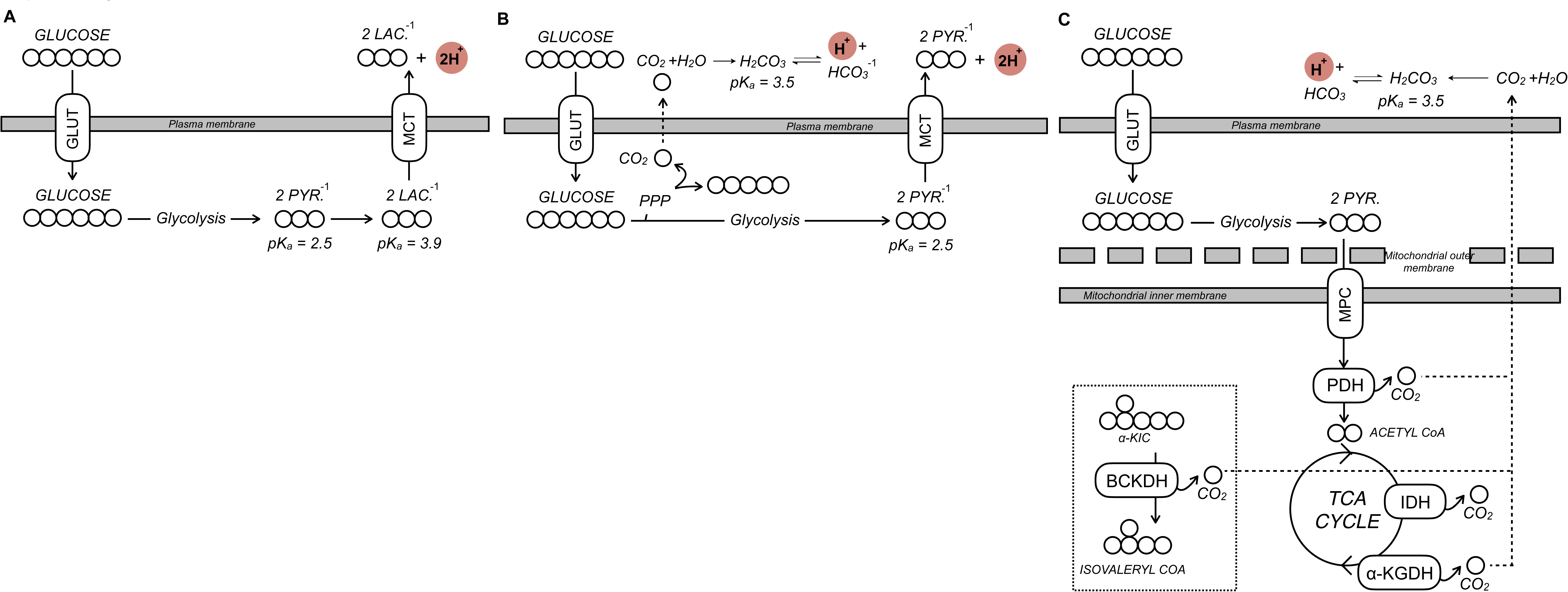
Metabolic reactions that generate net pH changes. (A) The main driver of extracellular acidification in most cell culture monolayer systems is lactate efflux. Uncharged glucose is taken up by the cell and converted to anionic pyruvate via glycolysis. Subsequent fermentation to lactate and efflux via monocarboxylate transporters causes a net pH change in the experimental medium that is detected by the XF Analyzer. GLUT, glucose transporter; MCT, monocarboxylate transporter; PYR., pyruvate; LAC., lactate. (B) Other reactions can cause a net pH change in the experimental medium, such as the efflux of glucose-derived carbon as anionic pyruvate or CO_2_ evolved from the oxidative pentose phosphate pathway (PPP) that ultimately forms bicarbonate and H^+^. (C) CO_2_ evolution by mitochondrial dehydrogenases also generates bicarbonate that can be detected by ECAR. CO2 is evolved from various mitochondrial enzymes including isocitrate dehydrogenase (IDH) and the three oxoacid dehydrogenases: pyruvate dehydrogenase (PDH), α-ketoglutarate dehydrogenase (α-KGDH), and the branched chain keto acid dehydrogenase (BCKDH). α-KIC, α-ketoisocaproic acid, the keto acid derivative of leucine.

**Expanded View Figure 2.**
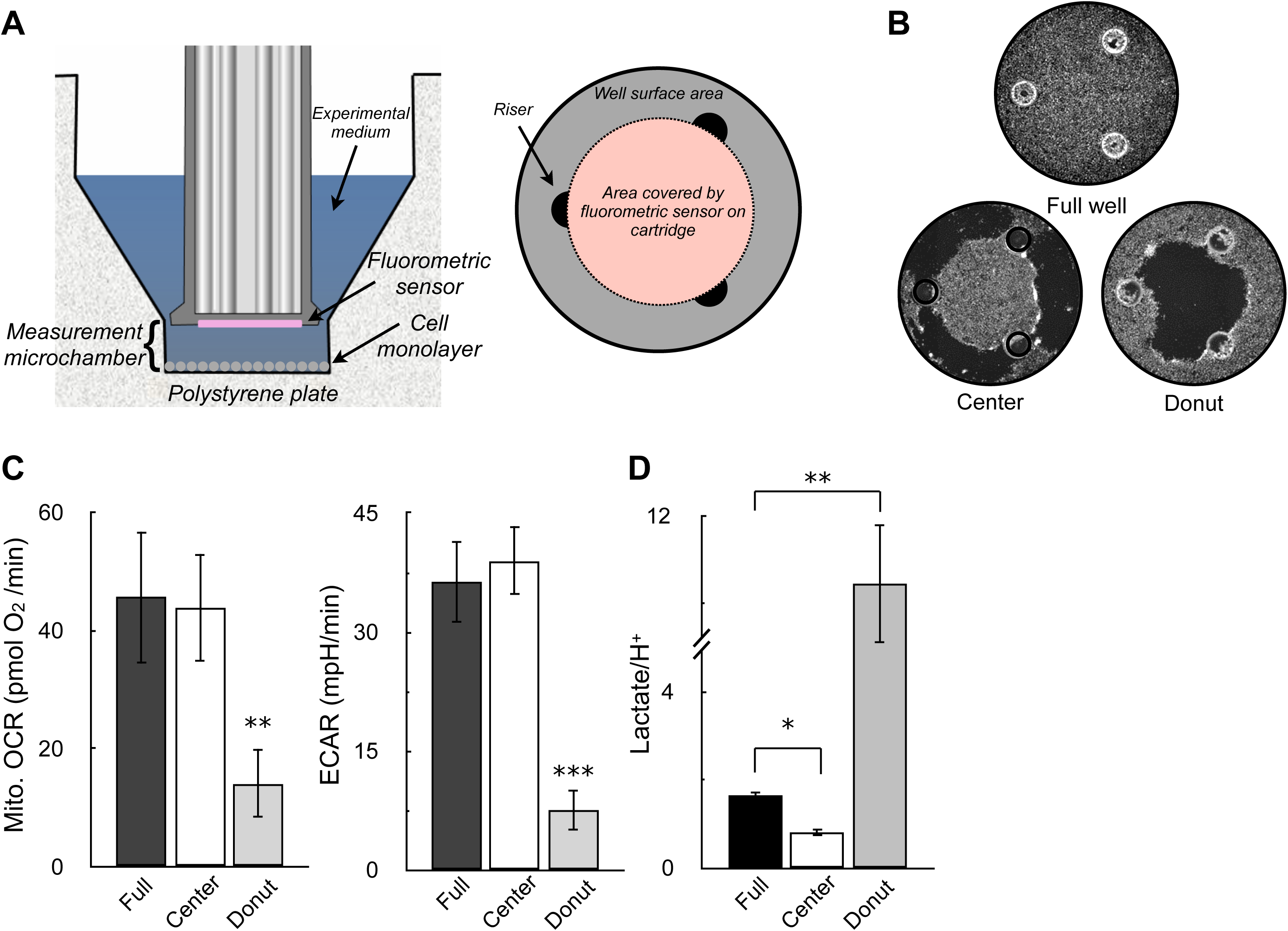
Sensor coverage of the well plate can explain why lactate:H^+^ > 1. All data are mean ± S.E.M. unless otherwise specified. *, p<0.05; **, p<0.01 (A) (*Left*) Cutaway image of Seahorse XF microplate and measurement sensor depicting location of the measurement microchamber, cell monolayer, and fluorometric sensor attached to the measurement cartridge. (*Right*) Birds-eye view diagram of the XF microplate well, where the confluent monolayer in drawn in grey, the three plate risers maintaining the uniform volume of the measurement microchamber are drawn in black, and the area covered by the fluorometric sensor is given in pink. (B) Representative images of wells assayed with an uninterrupted monolayer of cells (“Full well”), wells with cells scraped off the outer rim (“Center”), and wells with cells scraped off the inner portion of the well (“Donut”). (C) OCR and ECAR in A549 cells for the scraping conditions described in (B). (n=4 biological replicates) (D) The ratio of lactate (measured by enzymatic assay) to H^+^ (measured by the XF Analyzer) for the conditions described in (A & B). (n=4 biological replicates)

**Expanded View Figure 3.**
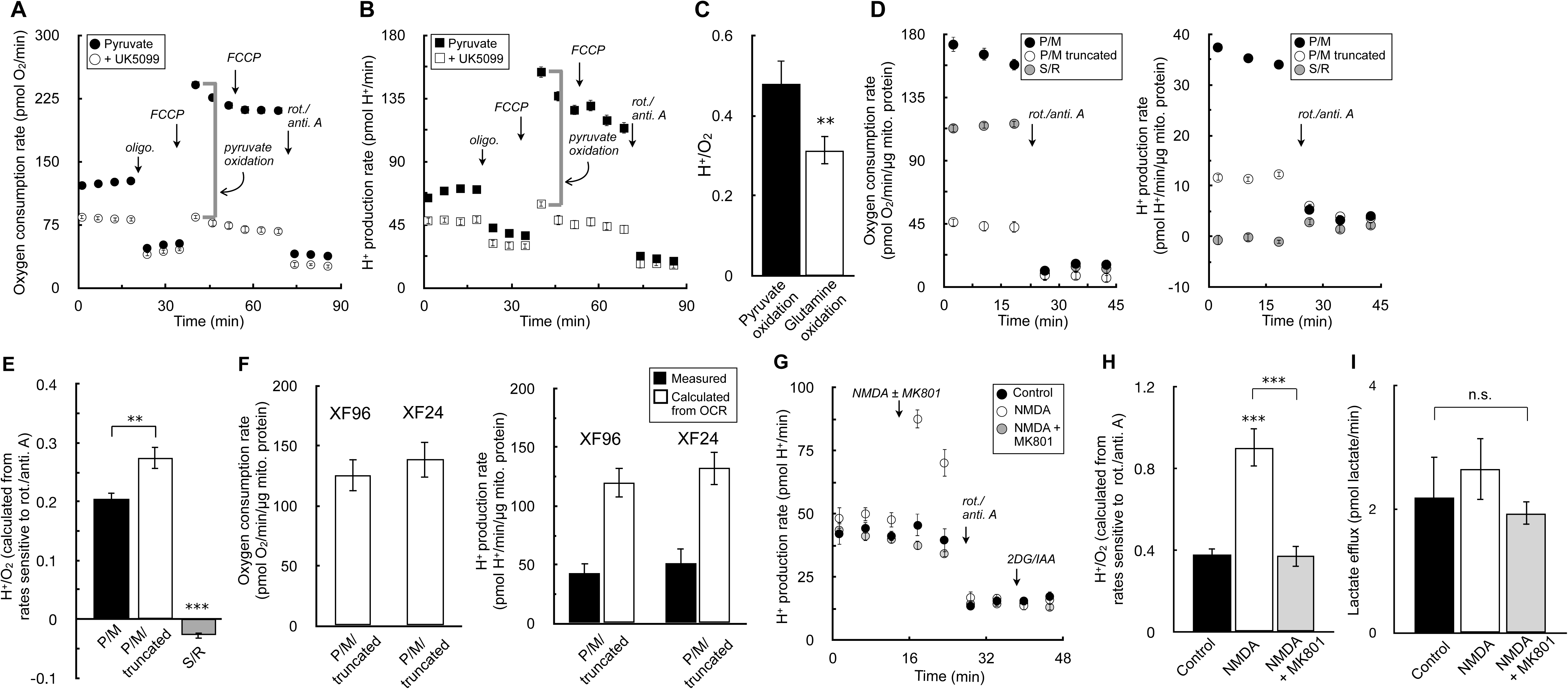
Cellular substrate preference and non-glycolytic organic acid efflux can affect H^+^:O_2_ ratios. All data are mean ± S.E.M. unless otherwise specified. *, p<0.05; **, p<0.01, ***, p<0.001 (A) Sample oxygen consumption kinetic trace for HepG2 cells offered 5 mM pyruvate, 5 mM 2-deoxyglucose (2-DG), and 50 μM iodoacetate (IAA) in the experimental medium. Where indicated, cells were treated with 1 μM UK5099 for 20 min prior to measurements. (n=10 technical replicates) (B) Sample H^+^ production rates for the conditions as in (A). (n=10 technical replicates) (C) The H^+^:O_2_ ratio is given for pyruvate and glutamine oxidation. Rates of H^+^ and O_2_ production for pyruvate oxidation were calculated as the UK5099-sensitive, maximal rates of respiration & acidification as indicated by the grey brackets in (A & B). Rates for glutamine oxidation were calculated similarly except 5 mM glutamine was offered instead pyruvate in the experimental medium, and the glutaminase inhibitor CB-839 was used instead of UK5099. (n=4 biological replicates) (D) Sample kinetic traces of oxygen consumption (*left*) and H^+^ production (*right*) for rat heart mitochondria offered 10 mM pyruvate with 1 mM malate and 2 mM dichloroacetate (P/M), a substrate and inhibitor mix to run a truncated TCA cycle consisting of P/M supplemented with 60 μM fluorocitrate (to block isocitrate dehydrogenase), 2 mM malonate (to block succinate dehydrogenase), and 1 mM aminooxyacetate (to block transaminase activity) (P/M truncated), or 10 mM succinate with 2 μM rotenone (S/R). All respiration measurements were made in the presence of 2 μM oligomycin and 4 μM FCCP, and background rates were calculated in response to 200 nM rotenone and 1 μM antimycin A. (n=20 technical replicates) (E) The H^+^:O_2_ ratio is calculated for each condition as in (D). (n=6 biological replicates) (F) (*Left*) Oxygen consumption rates in rat heart mitochondria offered conditions as in (D) for P/M truncated assayed in both the XF24 and XF96 assay platforms. (n=5 biological replicates) (*Right*) Measured H^+^ production rates for conditions as before (closed bars), compared to the theoretically predicted value assuming every mol of acid is stoichiometrically read by the instrument’s H^+^ sensor (open bars). (G) Primary cortical neurons are offered 5 mM β-hydroxybutyrate in aCSF medium supplemented with 5 mM 2-DG, and 50 μM IAA. Where indicated, cells were acutely treated with 100 μM NMDA or NMDA plus the NMDA receptor inhibitor MK-801 (10 μM). (n=10 technical replicates) (H) The H^+^:O_2_ ratio is calculated for each condition as in (F). (n=4 biological replicates) (I) The lactate efflux rate as measured by enzymatic assay for neurons treated for 20 min with the conditions as in (F). (n=3 biological replicates)

**Expanded View Figure 4.**
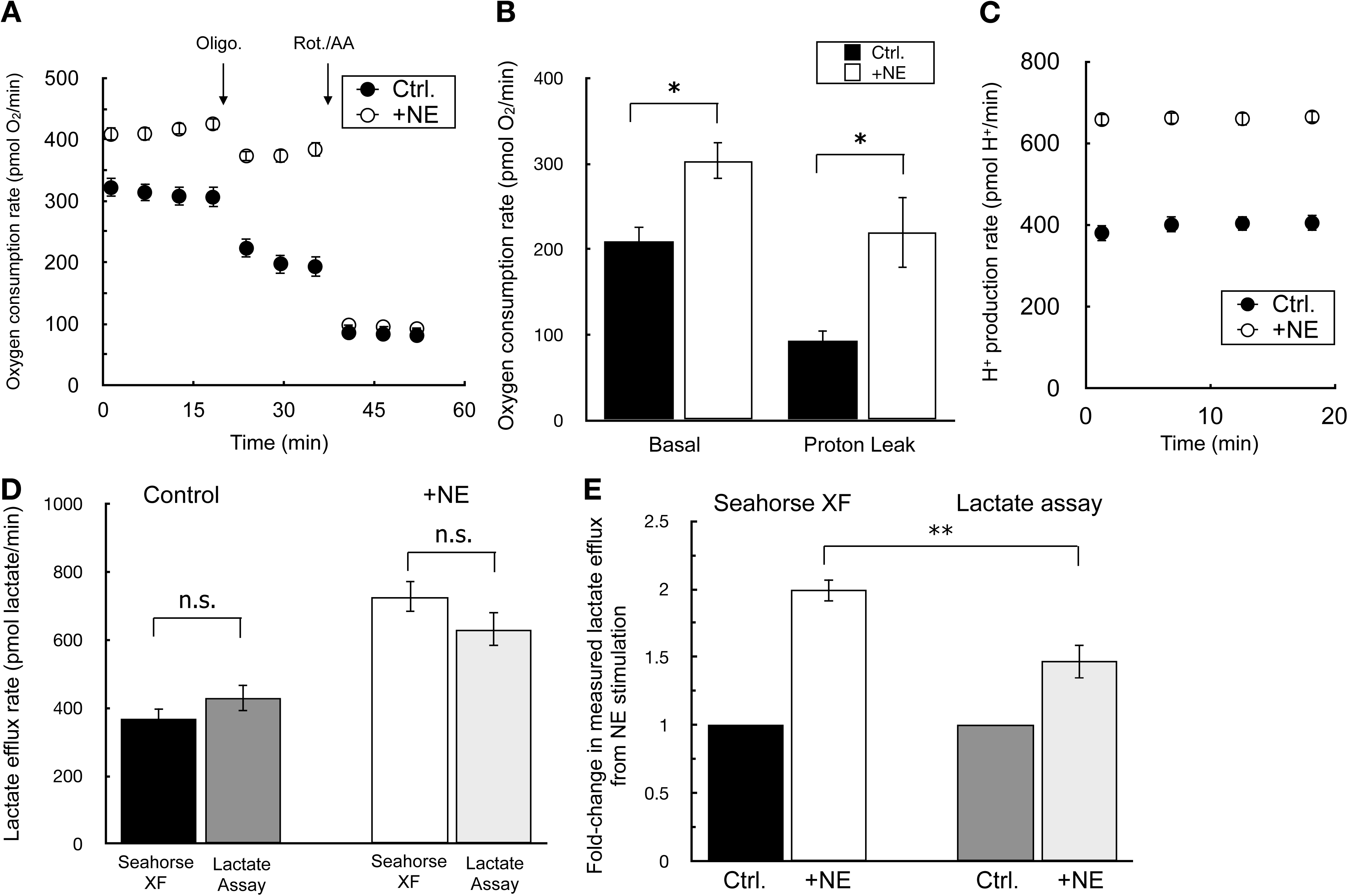
Changes in proportion of organic acid efflux during cell activation. All data are mean ± S.E.M. unless otherwise specified. *, p<0.05; **, p<0.01, ***, p<0.001 (A) Sample oxygen consumption kinetic trace for iBAT cells offered 10 mM glucose, 2 mM glutamine, and 2 mM pyruvate in the experimental medium. Where indicated, cells were treated with 2 μM oligomycin or 200 nM rotenone with 1 μM antimycin A. NE = 5μM norepinephrine. (n=10 technical replicates) (B) Sample kinetic trace of the H^+^ production rate as in (A). (n=10 technical replicates) (C) Rates of basal mitochondrial respiration rates and respiration associated with proton leak for conditions as in (A). (n=3 biological replicates) (D) Lactate efflux rates calculated by either the Seahorse XF Analyzer or an enzymatic lactate assay as described in the manuscript methods. NE, as before. (n=3 biological replicates) (E) Fold-change in lactate efflux in response to norepinephrine as measured by the Seahorse XF Analyzer or an enzymatic lactate assay. NE, as before. (n=3 biological replicates)

**Expanded View Figure 5.**
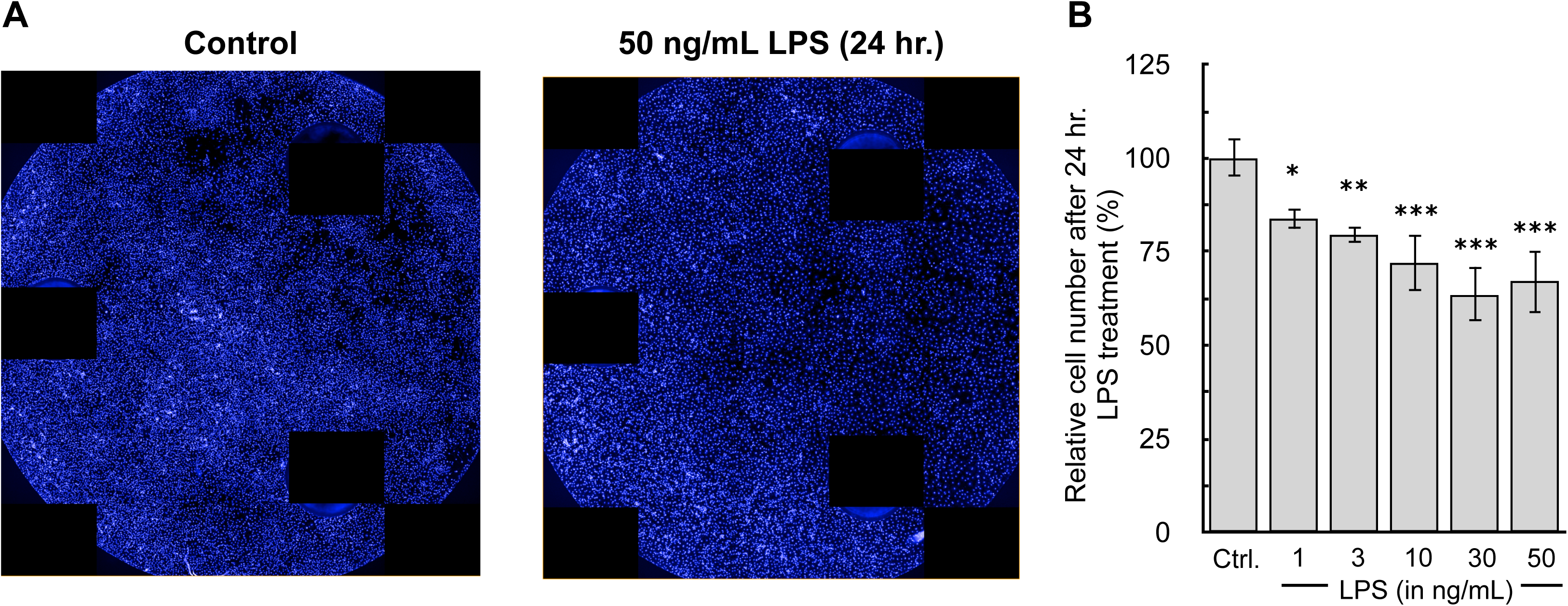
Images of well plate after 24 hr. LPS treatment. All data are mean ± S.E.M. unless otherwise specified. *, p<0.05; **, p<0.01, ***, p<0.001 (A) BMDMs were plated at 5×10^4^ cells/well treated with 50 ng/mL LPS for 24 hr. in Seahorse XF96 well plates. (A) After the assay, cells were fixed overnight with 2% (v/v) paraformaldehyde, stained with Hoescht 33342, and images were captured using the Perkin Elmer Operetta. (B) Quantification of cell counts after 24 hr. treatment with LPS. (n=4 biological replicates)

**Appendix Table 1:** **Lactate:H^+^ values used for composite factor**

| **<u>Cell Type</u>** | **<u>Lactate:H<sup>+</sup></u>** |
| --- | --- |
| 3T3-L1 | 1.73 |
| A431 | 1.28 |
| A549 | 1.60 |
| BAEC | 1.58 |
| BMDM | 1.58 |
| BT474 | 1.51 |
| C2C12 | 1.46 |
| Cortical neuron | 1.45 |
| H460 | 1.58 |
| HCT116 | 1.43 |
| HepG2 | 1.36 |
| HUVEC | 1.55 |
| Jurkat | 1.54 |
| MCF10A | 1.64 |
| MDA-MB231 | 1.61 |
| PC12 | 1.74 |
| Raw 264.7 | 1.45 |
| **Average** | **1.53** |
| **St. Dev.** | **0.12** |

**Appendix Table 2:** **H^+^:O_2_ values used for composite factor**

| **<u>Cell Type</u>** | **<u>H<sup>+</sup>/O<sub>2</sub></u>** |
| --- | --- |
| 3T3-L1 | 0.40 |
| A431 | 0.35 |
| A549 | 0.39 |
| BAEC | 0.34 |
| BMDM | 0.47 |
| C2C12 | 0.33 |
| Cortical astrocyte | 0.43 |
| Cortical neuron | 0.49 |
| H460 | 0.40 |
| HCT116 | 0.35 |
| HepG2 | 0.36 |
| HUVEC | 0.45 |
| INS-1 | 0.34 |
| Jurkat | 0.38 |
| MCF10a | 0.31 |
| MCF7 | 0.35 |
| MDA-MB231 | 0.38 |
| NRVM | 0.33 |
| PC12 | 0.41 |
| RAW 264.7 | 0.28 |
| Splenocytes | 0.35 |
| **AVERAGE** | **0.38** |
| **St. Dev.** | **0.05** |

